# From sediment to coral symbionts: persistent spatial structure across reef microbiomes

**DOI:** 10.1101/2025.06.23.661127

**Authors:** C. B. Scott, M. R. Collins, E. Nixon, A. K. Huzar, D. M. Flores, M. V. Matz, K. L. Black

## Abstract

Environmental context can structure microbial communities across space and time, but it remains unclear whether these patterns extend across free-living and host-associated communities or how they interact with host genetic background. We explored these relationships in reef sediments and the reef-building coral *Orbicella faveolata* in St. Croix, US Virgin Islands. We first repeatedly sampled sediments and individual coral colonies at two reefs over three weeks to compare microbial and algal symbiont (Symbiodiniaceae) communities between reefs and through time. Sediment microbial communities differed strongly between reefs and changed over the three-week period. Their functional composition showed similar patterns, indicating that taxonomic changes were accompanied by shifts in functional potential. In contrast, Symbiodiniaceae communities in both reef sediments and repeatedly sampled coral colonies differed strongly between reefs but remained largely stable through time. We then expanded our analysis across St. Croix to test whether host genetic differences could explain spatial variation in coral-associated Symbiodiniaceae. Symbiodiniaceae composition varied strongly among sites and with depth but was not associated with the two known genetic groups of *O. faveolata*. Together, these results show that components of the reef microbiome differ in their temporal dynamics but share strong spatial structure. While sediment microbial communities changed over just three weeks, Symbiodiniaceae communities were comparatively stable through time, and spatial differences among coral-associated Symbiodiniaceae persisted across the island regardless of host genetic background. Thus, reef site emerges as a strong and persistent source of variation across both free-living and host-associated components of the reef microbiome.

## Introduction

Coral reefs host a diverse array of microorganisms living in seawater, sediment, and coral hosts. These communities are highly dynamic, varying across space and time in response to shifting environmental conditions (Frade et al., 2020; Weber & Apprill, 2020). Seawater microbial communities in particular have been proposed as indicators of coral reef health, given their sensitivity to water quality and climate stress (Glasl et al., 2017; McDole et al., 2012). Although most scleractinian corals acquire at least some microbial partners from the surrounding environment, the communities that establish within hosts do not necessarily reflect the composition of that environment. Direct comparisons show that host-associated microbiomes are less closely related to environmental conditions than free-living communities, whether assessed taxonomically or functionally (Glasl et al., 2019, 2020; Zhao et al., 2023). These differences persist across reef sites, with coral and seawater microbiomes each varying by site while remaining compositionally distinct from one another (Lima et al., 2023).

Yet most evidence for environmental influences on coral-associated microbiomes comes from comparisons with the water column, leaving the reef sediment comparatively understudied. Like other reef microbial compartments, sediment communities vary substantially across depth, season, and reef location (Schöttner et al., 2011), and their close association with coral colonies creates a potential pathway for this environmental variation to influence host-associated microbiomes. Sediment immediately adjacent to coral colonies harbors distinct, more metabolically active bacterial assemblages than sediment further away (Pin et al., 2026), and coral mucus shares roughly twice as many bacterial taxa with nearby sediment as with the overlying seawater (Glasl et al., 2017). Differences in sediment bacterial communities have also been associated with differences in coral larval survival and settlement, suggesting that variation in the sediment microbiome may have consequences for coral recruitment (Serrano et al., 2024). Together, these observations suggest that sediment represents an important environmental microbial reservoir for corals, yet its contribution to the microbial communities established within hosts remains poorly understood.

This uncertainty may be particularly consequential for sediment-associated Symbiodiniaceae, the algal endosymbionts of scleractinian corals. Symbiodiniaceae are negatively buoyant and accumulate in reef sediments after release from hosts or other free-living sources, making sediment an important potential reservoir of environmental symbionts (Littman et al., 2008). The composition of these sediment-associated communities varies through time (Quigley et al., 2022) and over small spatial scales within individual reefs (Fujise et al., 2021), suggesting that corals may encounter distinct environmental symbiont pools across relatively short distances or timescales. Further, sediment exposure affects initial symbiont acquisition, suggesting that these environmental communities play a role in horizontal transmission, the primary route by which most coral species acquire their symbionts (Ali et al., 2019; Nitschke et al., 2016). Because established coral–symbiont associations tend to stabilize after an initial promiscuous period (Cumbo et al., 2013), the environmental pool encountered during transmission may have lasting consequences for which symbionts a colony ultimately hosts. Yet evidence for spatial and temporal variation in sediment-associated Symbiodiniaceae comes from only a handful of systems and remains largely disconnected from within-host dynamics (Bell & Quigley, 2025). Thus, variation in sediment-associated Symbiodiniaceae during coral spawning and recruitment could represent an important, yet poorly understood, source of variation in the symbiont communities that establish within coral hosts.

Beyond environmental variation, host genetic identity may also structure which symbiont communities are established within hosts. In hospite Symbiodiniaceae communities commonly vary among reefs and across environmental gradients, producing strong spatial structure in coral–symbiont associations (Denis et al., 2026; Howells et al., 2013; Kemp et al., 2015). At the same time, genomic studies are revealing widespread cryptic diversity within scleractinian corals, and these cryptic genetic groups frequently harbor distinct Symbiodiniaceae communities (Grupstra et al., 2024; Scott et al., 2025). Because this cryptic diversity is itself frequently structured across reef environments, spatial patterns in symbiont communities can be difficult to attribute to environmental availability or genetic differences among hosts.

The Caribbean reef-building coral *Orbicella faveolata* provides a well-studied system in which environmental conditions, local symbiont availability, and host genotype have all been implicated in structuring coral–algal associations. Symbiodiniaceae communities within *O. faveolata* vary across geographic and environmental gradients (Kemp et al., 2015; Parker et al., 2020; Rowan et al., 1997), while populations of common symbionts can remain strongly differentiated among reefs through time (Thornhill et al., 2009).

Studies incorporating host genetic variation have further demonstrated that these spatial patterns may reflect environmental conditions, host genetics, or both. In the closely related *O. annularis*, broad-scale Symbiodiniaceae biogeography was more strongly associated with environmental conditions than host population genetic structure (Kennedy et al., 2016). Conversely, host genet explained substantial variation in symbiont associations among individual *O. faveolata* colonies even after accounting for reef location (Manzello et al., 2019). Further, newly settled *O. faveolata* recruits tended to acquire symbiont communities according to their outplanting site rather than their parental lineage, implicating local reef conditions or differences in the available symbiont pool in initial symbiont establishment (Coffroth et al., 2022). Together, these studies demonstrate strong spatial structure in *O. faveolata* symbioses, but whether these site-specific associations reflect the local environmental symbiont pool, host genetic structure, or other features of the reef environment remains unresolved.

Here, we connected variation in the broader sediment microbiome to environmental and host-associated Symbiodiniaceae while accounting for cryptic host genetic structure in St. Croix, USVI. We combined repeated sampling of sediment and adult *O. faveolata* at two reefs with an island-wide survey spanning two cryptic host genetic groups distributed across eight reefs (Cabacungan et al., 2025). We conducted our temporal sampling surrounding the annual coral spawning event, coinciding with increasing sea surface temperatures, which we hypothesized would be the time of year most likely to induce short-term shifts in microbial communities. We asked whether (1) sediment microbial taxonomy and functional potential varied between reefs and through time, (2) environmental and host-associated Symbiodiniaceae showed similar spatial and temporal patterns, and (3) host genetic identity or local reef context better explained island-wide variation in coral-associated Symbiodiniaceae. Together, this design allowed us to test whether spatial patterns in the broader reef microbiome extend to coral-associated symbionts, or whether symbiont communities are instead structured by the genetic background of individual hosts.

## Methods

This study combined two sampling efforts—a spatial coral survey and a temporal coral-and- sediment survey—which together generated five molecular datasets from a 2022 field season in St. Croix, USVI (Supplemental Table 1). Both datasets collected *Orbicella faveolata* and/or surrounding reef benthos from 2-8 sampling sites (Figure 1a). All sampling was conducted in accordance with the U.S. Virgin Islands Department of Planning and Natural Resources (The Nature Conservancy Coral Restoration Permit DFW20052X).

**Figure 1.**
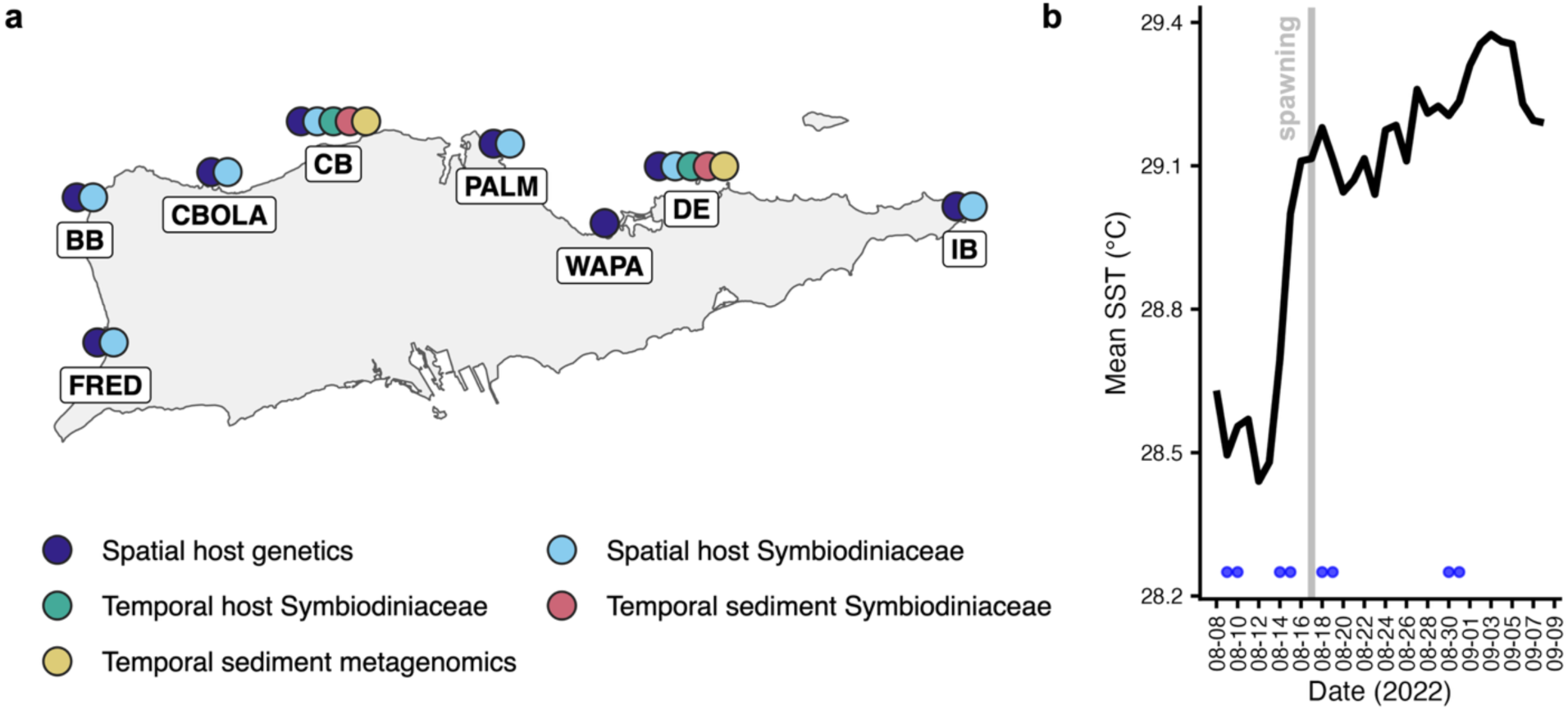
Spatial and temporal sampling design across St. Croix. (a) Locations and types of sampling conducted at eight reef sites. Color indicates the type of sampling conducted at each location (b) Mean sea-surface temperature (SST) across CB and DE during the 2022 sampling period. Blue points indicate sampling dates (CB was always sampled the day prior to DE), and the gray vertical line marks the date spawning was observed at DE for *O. faveolata*. Site abbreviations: BB, Butler Bay; CB, Cane Bay; CBOLA, Carambola; DE, Deep End; FRED, Frederiksted Pier; IB, Isaac Bay; PALM, The Palms; and WAPA, Water and Power Authority

### *O. faveolata* Spatial Survey

#### Sampling & laboratory methods

*O. faveolata* (Initial N=98) was sampled from 8 reef sites surrounding St. Croix, USVI in August 2022 (Figure 1). Adult colonies > 5cm in diameter were sampled via chiseling of a small portion from the colony base. For each sample, depth was recorded *in situ*. Tissue samples were stored in 100% ethanol at -20 °C until DNA was extracted.

Genomic DNA was isolated using a CTAB lysis buffer and organic DNA extraction (detailed in Supplemental File 1). Extraction was followed by purification using the Zymo Genomic DNA Clean & Concentrator kit (Zymo #D4067).

#### Host genotyping

Hosts were genotyped using 2bRAD, a restriction site-associated DNA sequencing method (Wang et al., 2012). The resulting 2bRAD libraries were sequenced at the Genomic Sequencing and Analysis Facility at the University of Texas at Austin on the Illumina NovaSeq SR100 platform. Sequencing was ultimately conducted in two batches, as only 47 individuals out of the 98 collected were successful after the initial sequencing effort (see Cabacungan et al., 2025). After repreparing libraries and resequencing failed samples, we retained a total of 66 individuals after the quality filtering subsequently outlined.

Raw sequences were processed using the Texas Advanced Computing Center (TACC). Raw reads were trimmed and deduplicated following the pipeline hosted at https://github.com/z0on/2bRAD_denovo. Reads were then mapped to a concatenated reference containing the *O. faveolata* host genome (Young et al., 2024) and *Symbiodinium* sp. (Shoguchi et al., 2018)*, Breviolum minitum* (Shoguchi et al., 2013), *Cladocopium* sp. (Shoguchi et al., 2018), and *Durusdinium trenchii* (Dougan et al., 2024) genomes using bowtie2 (Langmead & Salzberg, 2012) in local alignment mode. Alignments were filtered to a minimum mapping quality of 20 and sorted with SAMtools (H. Li et al., 2009). To isolate host-derived data, we removed reads mapping to Symbiodinacaeae genomes.

Genotype likelihoods were calculated in ANGSD (Korneliussen et al., 2014). We first ran an initial quality-assessment pass on all mapped, filtered bam files, requiring unique, non-duplicate reads, a minimum mapping quality of 20, a maximum depth of 1000, and data present in at least 60 individuals. This pass generated per-sample quality score distributions and depth and count statistics. Based on these metrics, we retained 77 individuals for downstream analysis.

Using this filtered sample set, we called genotype likelihoods and SNPs in ANGSD with the following filters: unique, non-duplicate reads only, a minimum mapping quality of 20, a minimum base quality of 25, exclusion of triallelic sites, exclusion of sites showing significant strand bias or heterozygosity bias (*p* < 10^-5^ for both), a minimum minor allele frequency of 0.05, and requiring genotype likelihoods in at least 75% of the 77 samples. This retained approximately 20,000 SNPs. Genotypes were compiled into a pairwise identity-by-state (IBS) genetic dissimilarity matrix for initial inspection. Hierarchical clustering of the genetic matrix was evaluated for correct alignment of technical replicates, and identification of clonal samples. After clone removal, samples were re-genotyped with ANGSD with the same filters outlined above, retaining a total of 66 samples.

We used the resulting IBS matrix to examine cryptic genetic structure among samples. This structure recapitulated the two major clusters previously identified by Cabacungan et al. (2025) based on the initial 47 samples. We assigned samples to these clusters via average-linkage hierarchical clustering of the IBS matrix, cutting the tree at a height of 0.3.

#### Symbiodiniaceae data generation

Amplicon sequencing libraries were constructed from the DNA extractions detailed above for genotyped *O. faveolata* samples. ITS2 amplification used primers from (Hume et al., 2018); SymVarF: CTACACGACGCTCTTCCGATCTGAATTGCAGAACTCCGTGAACC, SymVarR: CAGACGTGTGCTCTTCCGATCTCGGGTTCWCTTGTYTGACTTCATGC). All sequencing libraries were amplified with 20-30 PCR cycles, depending on band intensity. Negative controls (MilliQ water) were run with each set of library preparations and amplification was never observed. Libraries were pooled according to visually assessed gel band brightness. Pooled amplicon libraries were sequenced on the Illumina MiSeq v3 for PE300 reads.

Raw reads were processed through the Symportal framework (Hume et al., 2019), which performs quality control, chimera detection, and ITS2 profile assignment using its own reference database and defined intragenomic sequence variant-based framework. After read filtering and quality control we retained 80 samples. In total, 55 samples had both successful host genotyping and Symbiodiniaceae community assignments.

### Temporal Reef Survey

#### Sampling

Samples were collected on the north shore of St. Croix, USVI between August 9, 2022, and August 31, 2022. We collected *O. faveolata* and marine sediment samples at two reefs (Cane Bay, CB, and Deep End, DE) across 4-5 time points (Figure 1a). All samples were collected between 10AM and 2PM, except for sediment samples taken on the night of spawning at DE. In an attempt to increase our likelihood to detect microbial community turnover, temporal sampling coincided both with the annual coral spawning event and with an increase in mean sea surface temperature (meanSST; Figure 1b). Coral spawning was observed *in situ* for *O. faveolata* at DE on the night of August 16, 2022. Due to the distance between CB and DE and personnel limitations, we could not verify spawning in situ at CB. However, it is highly likely that *O. faveolata* spawning occurred at CB on August 16-17, as spawning was reported or predicted for the species during this window at sites as distant as the Florida Keys and the Bahamas (Hurtado-López et al., 2022; Williams et al., n.d.).

Four *O. faveolata* genotypes were tagged at CB and three were tagged at DE for repeated sampling. For each *O. faveolata* tissue sample, three polyps were scratched (two on opposing sides, one on top of the colony) using a 16G needle attached to a 20 mL Luer-lock syringe. The three polyps collected from each colony were combined during sampling into one colony-average measure.

At each site and sampling date, we collected three replicate samples from the upper 2 cm of sediment at each of three subsites, yielding nine sediment samples per site and date. Samples were collected in 2-mL cryovials and processed separately. One subsite was located within 5 m of the tagged coral colonies, while the other two were randomly selected along a cross-shelf gradient. We recorded depth at each subsite. After quality filtering, we retained 20 samples from CB and 7 from DE for ITS2 analysis and 27 from CB and 28 from DE for metagenomic analysis.

Samples were processed within two hours of collection at the Nature Conservancy’s Coral Innovation Hub, in St. Croix, USVI. Coral tissue was first rinsed in 100% EtOH, then fixed in 100% EtOH and stored at -20°C until DNA extraction. Sediment samples were centrifuged in a tabletop centrifuge for two minutes. Seawater was then removed from top of the vial and replaced with 100% EtOH and samples were stored at -20°C.

#### DNA extraction, WGS library preparation & read processing

Host tissue was extracted using the same method as the spatial survey detailed above. Sediment samples were processed using the Qiagen PowerSoil Pro Kit [catalog #47014]. We used 0.125g of sediment as the input for each extraction and added an additional 50 μL of Qiagen Buffer ATL to 750 μL of Qiagen Buffer C1 during the lysis step. We additionally prepared one CTAB and PowerSoil Kit extraction negative, using 2 μL MilliQ water as a blank input.

Low-coverage whole-genome shotgun (lcWGS) sequencing libraries were prepared for host and sediment samples utilizing unloaded Tn5 transposase (see detailed protocol in Supplemental File 2). Briefly, we assembled the tagmentase complex by loading the Tn5 transposase with custom designed adaptors that are complementary to the 19 bp consensus region on the Tn5 transposase as well as to the barcodes. The tagmentase complex was diluted to 50 ng/μl before being added to DNA (2.5 total ng). DNA was then amplified and barcoded in one PCR reaction. All libraries were amplified along with a PCR negative control, which did not show any amplification. We then sequenced libraries at the University of Texas Genomic Sequencing and Analysis Facility on the Illumina NovaSeq SP with a target of five million PE 100bp reads per sample.

Illumina adapters were trimmed from paired reads with Trim Galore using default settings and allowing the autodetection of adapters (https://github.com/FelixKrueger/TrimGalore). The remaining “mosaic” portions of adaptors, designed to achieve base call diversity at the beginning of reads, were trimmed using Cutadapt (Martin, 2011).

#### Taxonomic assignment of lcWGS data

SingleM was used to profile our shotgun metagenomes from the processed reads (Woodcroft et al., 2026). SingleM uses hidden markov models to identify single copy archaeal, bacterial, and cross-domain OTUs in metagenomic sequencing reads. This improves abundance estimation and enables the removal of eukaryotic host contamination from microbial community data. In brief, singleM was run on paired-end inputs, using the “smafa_naive_then_diamond” assignment method, and requiring a minimum taxon coverage of 0.1 in the consensus profile table. We then used the consensus taxonomic profile table from singleM for subsequent community-level analyses. Notably, singleM identified very few non-coral taxa for our host-associated samples, therefore these were omitted from subsequent analyses. SingleM did not identify any OTUs in negative controls for CTAB and PowerSoil extractions. After filtering and taxonomic assignment, 77 sediment samples were retained for subsequent analysis.

#### Functional annotation of lcWGS data

We co-assembled contigs from sediment samples using MEGAHIT (D. Li et al., 2015) with the ‘meta-large’ preset. Contigs were then classified as prokaryotic or eukaryotic in origin using Tiara (Karlicki et al., 2022), requiring a minimum contig length of 1000 bp and a classification probability of at least 0.65. Contigs classified as bacteria, archaea, or generic prokarya were treated as the prokaryotic set, and contigs classified as eukaryotic were treated as the eukaryotic set.

Genes were predicted separately for each contig set. Prokaryotic genes were predicted using Prodigal in metagenome mode (Hyatt et al., 2010). Eukaryotic genes were predicted using MetaEuk (Levy Karin et al., 2020), searching against the UniRef50 reference database. Predicted protein sequences from both sets were then combined for functional annotation using eggnog-mapper. DIAMOND was used as the search method and ‘mid-sensitive’ was used as a search mode against the eggNOG reference database.

To quantify gene abundance, prokaryotic and eukaryotic contigs were combined into a single reference and indexed with BWA (H. Li & Durbin, 2009). Sediment metagenomic reads were mapped against this combined reference using bwa mem with default settings. Alignments were sorted and indexed with SAMtools (H. Li et al., 2009). Gene-level read counts were generated using featureCounts (Liao et al., 2014) against predicted gene coordinates from Prodigal and MetaEuk and normalized by gene length to yield reads per kilobase (RPK). Genes were then converted to presence-absence, with presence defined as RPK ≥ 1.

#### Symbiodiniaceae ASV generation

Our lcWGS approach did not yield sufficient coverage of Symbiodiniaceae in either host or sediment samples for downstream analysis. Therefore, we additionally generated ITS2 amplicon sequencing libraries from these samples following the protocol outlined above for the spatial surveys. Samples were sequenced in a separate batch on the Illumina MiSeq v3 for PE100 reads.

Paired-end ITS2 amplicon reads from our marine sediment and through-time hosts were processed using DADA2. We used the DADA2 pipeline for this task rather than Symportal because environmental samples do not conform to the assumptions of the Symportal framework and we wanted to compare ASVs from paired sediment and host samples directly. Reads were quality-filtered and trimmed, removing 10 bp from each end and truncating to a fixed length of 100 bp per read, as quality sharply declined beyond this. Reads falling below Q10 were truncated at that position, and PhiX-derived reads were removed. Samples with fewer than 50 reads after filtering were excluded. Error rates were learned separately for forward and reverse reads and used to denoise each read direction with DADA2’s inference algorithm.

#### Taxonomic assignment of ASVs

Taxonomy was assigned to each final ASV via BLAST search against NCBI’s nucleotide database, retaining only the top-scoring hit(s) by e-value (ties retained). Full taxonomy for each hit was then retrieved from its accession number. ASVs not assigned to family Symbiodiniaceae were excluded. Because some ASVs had multiple top-scoring hits, we assigned each a consensus taxonomy. Generic labels (e.g., “Symbiodinium sp.”) were dropped in favor of more specific hits when genus-level assignments agreed. Consensus taxonomy was then based on the most frequent species name among the remaining hits. If no species reached a majority, we used the most frequent genus. Similarly, if no genus reached consensus, the ASV was left unassigned below Symbiodiniaceae.

To reduce inflation of ASV richness from intragenomic ITS2 variation, ASVs were clustered by co-occurrence across samples. Pairwise Jaccard distances were calculated from ASV presence/absence across samples and used for average-linkage hierarchical clustering. Groups were defined by cutting the tree at a distance threshold of 0.05. Read counts were summed within each group per sample to produce a sample-by-group abundance table.

We then assigned a consensus taxonomy to each ASV group using the same approach. Groups with consistent genus-level assignments across member ASVs were labeled with the most common non-generic (species- or strain-level) label available. Groups with genus consensus but with no majority species assignment were labeled Genus sp. and groups lacking genus-level consensus were left unassigned beyond the family level.

### Statistical Analysis

We conducted all statistical analyses in R (version 4.5.2). Although each molecular dataset was analyzed separately, we used a common framework to test variation in community composition across space and time. We first describe analyses of spatial host-associated Symbiodiniaceae, followed by analyses of the temporal sediment and coral datasets and KO-level functional associations. For temporal analyses, sampling date was expressed as days relative to the observed *O. faveolata* spawning event, with negative values representing samples collected before spawning and day zero corresponding to August 16, 2022.

#### Spatial survey host-associated Symbiodiniaceae

We converted SymPortal ITS2 type-profile abundances to relative abundance, calculated Bray– Curtis dissimilarities among samples and used the term-wise PERMANOVA model ‘profile distance ∼ site * depth * host genetic group’ with 999 permutations.

Using site-level environmental data from Cabacungan et al. (2025), we next tested whether environmentally similar reefs supported similar host-associated Symbiodiniaceae communities. We represented Symbiodiniaceae composition by the first three axes of a PCoA of Bray-Curtis dissimilarities and calculated the centroid of each reef site in this ordination space. We generated corresponding Euclidean environmental distances between sites and compared the two configurations using a Procrustes rotation test with 999 permutations.

Because environmental conditions were measured at the site level, colonies within a site were not independent replicates of those conditions. We therefore jointly analyzed the first three Symbiodiniaceae MDS axes using two multivariate Bayesian Gaussian models. Both included depth and host genetic group as fixed effects. The first accounted for clustering among colonies from the same site using an independent random intercept for site. The second allowed site effects to covary according to environmental similarity. Comparing these models allowed us to test whether environmental similarity explained variation among sites beyond that captured by site identity alone. We fit both models with estimated residual correlations among axes. We assigned Normal(0, 1) priors to regression coefficients and assigned an LKJ(2) prior to the residual correlation matrix. We fit all models in brms (Bürkner, 2021) using the CmdStan backend, four chains of 6,000 iterations each and ‘adapt_delta = 0.99’. Posterior predictive checks and convergence diagnostics (Rhat <1.01, ESS > 1000) confirmed adequate fit across all models.

#### Temporal survey community analyses

For each temporal dataset, we calculated pairwise compositional distances between samples and tested the effects of the relevant spatial and temporal predictors using term-wise permutational multivariate analysis of variance (PERMANOVA; 999 permutations). For sediment microbial taxonomy, we calculated weighted UniFrac distances from the SingleM taxon-coverage matrix. Because SingleM provided taxonomic lineages rather than a phylogenetic tree with estimated branch lengths, we constructed a hierarchical tree from the reported lineage strings and assigned unit length to each taxonomic step. For sediment functional composition, we linked genes to KEGG Ortholog (KO) terms using the eggNOG-mapper annotations; genes assigned to multiple KOs contributed to each listed term. We classified a KO as present in a sample when at least one gene assigned to it was detected and calculated Jaccard distances from the resulting sample-by-KO presence–absence matrix. For sediment and temporally sampled host-associated Symbiodiniaceae, we centered-log-ratio transformed ASV-group abundances after adding a pseudocount of one and calculated Euclidean distances. We retained ASV groups detected in at least two samples and samples containing at least two ASV groups.

For sediment taxonomic and functional composition, PERMANOVA models had the form ‘compositional distance ∼ site * (sampling date + depth)’ The sediment-associated Symbiodiniaceae model had the form ‘compositional distance ∼ site * depth + site * sampling date’, and the model for host-associated Symbiodiniaceae sampled through time had the form ‘compositional distance ∼ site * sampling date’. We also tested concordance between sediment taxonomic and functional community patterns using a Procrustes randomization test of the weighted UniFrac and KO Jaccard distance matrices.

For each temporal dataset, we complemented the PERMANOVA with a multivariate Bayesian Gaussian model that accounted for repeated observations. We included sampling subsite as a random intercept for sediment samples and colony identity as a random intercept for repeatedly sampled corals. We represented each sample by its scores on the first three principal-coordinate axes of the corresponding distance matrix and analyzed the axes jointly. All models included site, sampling date and their interaction as fixed effects. We included sampling subsite as a random intercept for the sediment datasets and coral colony genotype as a random intercept for the repeatedly sampled hosts, giving the general model structure ‘community principal component ∼ site * sampling date + (1 | sampling unit)’. Models were fit with the Bayesian settings described above.

#### KO-level association analysis

To identify individual functions associated with site and sampling date, we fit a separate binomial generalized linear model for each KO that varied among samples, using the model ‘KO presence ∼ site * sampling date’. We adjusted Wald-test *p* values for multiple comparisons within each model term using the Benjamini–Hochberg false-discovery-rate procedure and considered KOs with an adjusted *p* < 0.05 significant.

#### AI-assisted functional interpretation

Significant KEGG Orthologs (KOs) identified by the GLM were interpreted using OpenAI Codex (Sol 5.6, accessed August 2026). The model was provided with the KO-level statistical results and eggNOG-mapper annotations, including GO terms, COG categories, KEGG pathways/modules, PFAM domains, CAZy assignments, and broad taxonomic annotations associated with the KO terms. It was instructed to group KOs into broad biological processes relevant to coral reefs, marine benthos, or marine environments, while prioritizing general biochemical function when reef-specific and broader interpretations differed. KOs representing housekeeping functions or insufficiently supported ecological assignments were retained as general or unresolved. Alternative KO assignments associated with the same annotated gene were collapsed into a single annotation evidence unit when evaluating support for a functional theme. The model assisted with literature discovery and synthesis but did not process the sequence data or perform the original differential-abundance GLM. KO definitions and key interpretations were checked against KEGG records, and supporting ecological interpretations were evaluated using the peer-reviewed literature. All final classifications and conclusions were reviewed by the authors. A list of KO to functional category translations and supporting logic can be found in Supplemental File 3.

## Results

### Sediment metacommunity taxonomy and gene families vary between reef sites and through time

First, we modeled how overall sediment microbial taxonomic composition varied between reef sites, among depths within reefs and through time. PERMANOVA indicated that reef site (CB versus DE) explained the largest proportion of variation in taxonomic composition (*R*² = 0.0759, *p* = 0.004), followed by the interaction between site and sampling day (*R*² = 0.0441, *p* = 0.028) and sampling day (*R*² = 0.0426, *p* = 0.030). Neither depth nor its interaction with site was significant (Supplemental Table 2).

We complemented this analysis with a Bayesian hierarchical model that accounted for repeated sampling of reef subsites through time, which largely recapitulated these results. The model supported a difference between sites along MDS2 (β = 0.07, CrI_95%_ = [0.04, 0.10]) and a weakly positive site-by-sampling day interaction along MDS1 (β = 0.01, CrI_95%_ = [0.00, 0.02]), reaffirming that temporal change along this axis differed between CB and DE. The remaining fixed-effect credible intervals overlapped zero (Figure 2a; Supplemental Tables 3-4).

**Figure 2.**
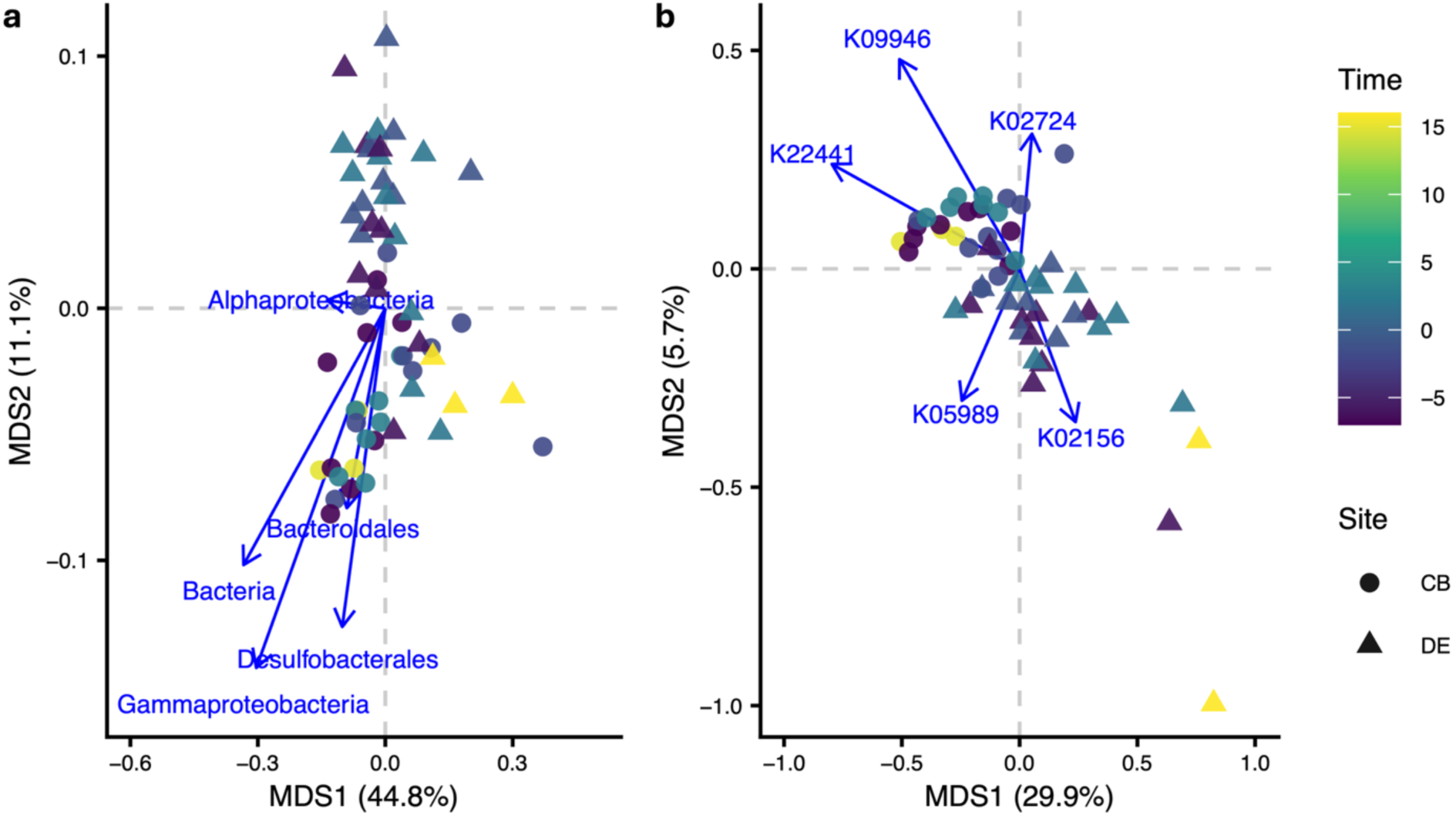
Site- and temporal variation in the taxonomic and functional composition of sediment microbial communities. Principal coordinates analyses of (a) taxonomic composition based on weighted UniFrac distances and (b) KEGG Ortholog (KO) composition based on Jaccard distances of KO presence–absence. Points represent individual sediment samples, with color indicating sampling time in days relative to observed *O. faveolata* spawning and shape indicating reef site (CB, Cane Bay; DE, Deep End). Blue arrows show selected taxa or KOs with strong, directionally distinct loadings on the displayed axes. Percentages indicate the variation represented by each axis.

Taxonomic and functional community configurations were strongly correlated (Procrustes sum of squares = 0.413, *r* = 0.7662, *p* = 0.001). PERMANOVA analysis indicated that functional composition also varied by reef site (*R*² = 0.1314, *F* = 8.55, *p* = 0.001), the interaction between site and sampling day (*R*² = 0.0407, *F* = 2.65, *p* = 0.014) and the interaction between site and depth (*R*² = 0.0391, *F* = 2.54, *p* = 0.017). Sampling day and depth alone were not significant (Supplemental Table 5). Functional composition was also more variable within DE than within CB, with greater mean distance to the group median at DE (0.120) than at CB (0.076; *F* = 17.11, *p* < 0.001).

A Bayesian hierarchical model supported the PERMANOVA results while accounting for repeated sampling of reef subsites. Functional composition differed between sites along MDS1 (β = 0.42, CrI_95%_ = [0.23, 0.62]), and the site-by-sampling day interaction was also positive along this axis (β = 0.03, CrI_95%_ = [0.01, 0.06]). This interaction supported that temporal change in functional composition differed between CB and DE. Credible intervals for the remaining fixed effects overlapped zero (Figure 2b; Supplemental Tables 6-7).

To identify the functions contributing to site-level differences in functional composition, we fit binomial models to the presence or absence of each KO. After Benjamini–Hochberg correction, 205 KOs differed significantly between sites (adjusted *p* < 0.05), with 202 occurring more frequently at CB and only three occurring more frequently at DE (Supplemental File 3). After excluding general and unresolved annotations and collapsing alternative KO assignments associated with the same gene, the most strongly represented CB-associated themes included eukaryotic cellular regulation and maintenance, stress tolerance and genome plasticity, cyanobacterial or oxygenic phototrophy, environmental sensing and regulatory flexibility, cell-envelope and biotic-interaction functions, aromatic-compound transformation and organic-matter processing. The three DE-associated KOs were assigned to eukaryotic lipid or redox metabolism, eukaryotic RNA silencing and genome defense, and Rieske oxygenase or xenobiotic transformation (Figure 3). No individual KOs differed significantly through time after multiple-testing correction.

**Figure 3.**
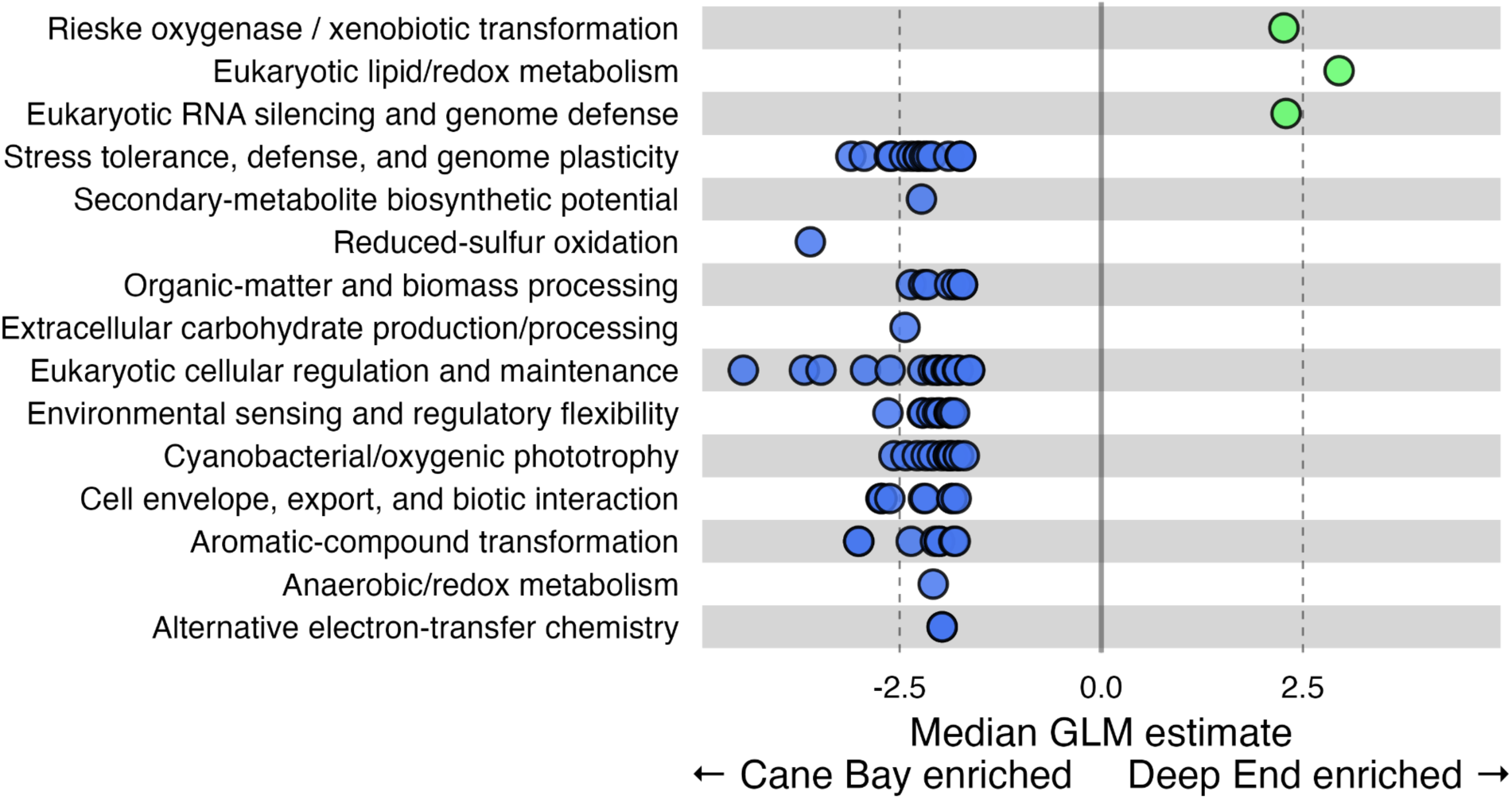
Site-associated differences in the functional potential of sediment microbial communities. Points represent KEGG Orthologs (KOs) whose occurrence differed significantly between Cane Bay and Deep End in binomial generalized linear models (Benjamini–Hochberg-adjusted *p* < 0.05) grouped by their assigned functional theme. Position indicates the site-effect estimate for KOs within each theme, with negative values indicating enrichment at Cane Bay (CB) and positive values indicating enrichment at Deep End (DE). General, unresolved and housekeeping functions are not shown.

### Sediment Symbiodiniaceae communities vary between reef sites and through time

Sediment-associated Symbiodiniaceae composition differed strongly between reef sites, which explained the majority of variation among samples (PERMANOVA; *R*² = 0.4004, *F* = 19.26, *p* = 0.001). The interaction between site and depth was also significant (*R*² = 0.0574, *F* = 2.76, *p* = 0.031), whereas depth alone was not significant. Neither sampling day nor its interaction with site was significant, indicating that sediment-associated Symbiodiniaceae composition remained largely stable over the sampling period (Supplemental Table 8; Figure 4a).

**Figure 4.**
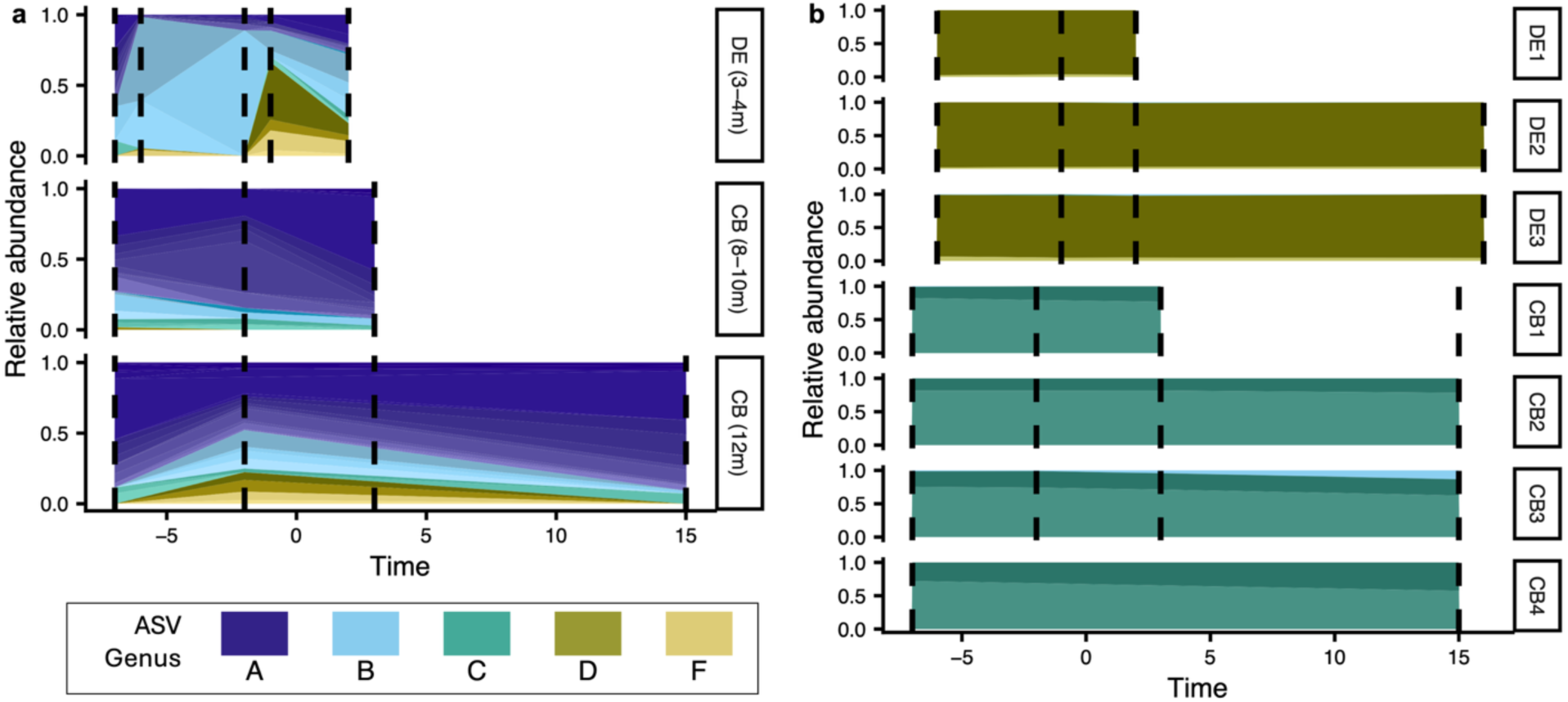
Temporal composition of Symbiodiniaceae ASV groups in sediment and coral hosts. Relative abundances of Symbiodiniaceae ASV groups in (a) sediment samples grouped by site and depth and (b) repeatedly sampled *O. faveolata* colonies. Sediment values represent mean relative abundance across replicate samples within each site–depth category and sampling date. Coral facets represent individual tagged colonies at Deep End (DE1–DE3; 3-4m) and Cane Bay (CB1–CB4; 12m). Time is expressed as days relative to observed spawning, and dashed vertical lines indicate sampling dates. Colors identify Symbiodiniaceae genera (A, *Symbiodinium*; B, *Breviolum*; C, *Cladocopium*; D, *Durusdinium*; F, *Fugacium*), with different shades representing distinct co-occurrence-defined ASV groups within each genus. Only ASV groups reaching at least 5% relative abundance in one or more samples are shown.

Although the PERMANOVA did not identify an overall effect of sampling day, the Bayesian hierarchical model supported a site-specific temporal pattern along MDS3. The site-by-sampling day interaction was positive along this axis (β = 0.47, CrI_95%_ = [0.10, 0.84]), indicating that temporal change in sediment-associated Symbiodiniaceae composition differed between CB and DE. Inspection of the ordination loadings indicated that MDS3 was driven primarily by the presence of uncultured *Symbiodinum*- and/or absence of *Breviolum*-associated ASV groups (Supplemental Figure 1). The model also supported a strong difference between sites along MDS1 (β = 2.41, CrI_95%_ = [1.71, 3.00]), consistent with the dominant site effect identified by PERMANOVA. Credible intervals for the remaining fixed effects overlapped zero. Finally, subsite-level variation was substantial along MDS2 (SD = 1.35, CrI_95%_ = [0.84, 2.09]) and MDS3 (SD = 0.92, CrI_95%_ = [0.41, 1.57]), consistent with persistent fine-scale spatial heterogeneity within reefs (Supplemental Tables 9-10).

### Reef site, not time or host genetics, predominantly structures within-host Symbiodiniaceae community

Conversely, Symbiodiniaceae communities in *O. faveolata* colonies tracked through time differed consistently between sites but showed no temporal change (Figure 4b). PERMANOVA found that site explained almost all variation in community composition (*R*² = 0.8581, *F* = 162.22, *p* = 0.001). Colony identity, sampling day and the interaction between site and sampling day were not significant (Supplemental Table 11).

The Bayesian hierarchical model accounted for repeated sampling of colonies and recovered a strong difference between sites along MDS1 (β = 2.02, CrI_95%_ = [1.70, 2.29]). This site contrast was large relative to variation among colonies on the same axis (SD = 0.10, CrI_95%_ = [0.00, 0.35]). Credible intervals for sampling day and the site-by-sampling day interaction widely overlapped zero on all three axes, providing no evidence of temporal change within either reef (Supplemental Tables 12-13).

Given the strong and temporally persistent differences between CB and DE, we next expanded our spatial scale to ask whether variation in host-associated Symbiodiniaceae was related to host genetic group or local reef conditions (Figure 5a,b). Across an island-wide survey, host-associated Symbiodiniaceae ITS2 type-profile composition differed strongly among reef sites, with site explaining the largest amount of variation among colonies (*R*² = 0.3119, *F* = 3.84, *p* = 0.001; Figure 5c,d). The interaction between site and depth was also significant in explaining variation (*R*² = 0.0964, *F* = 3.56, *p* = 0.006), though is likely driven by the comparatively large depth variation at CB and CBOLA (Supplemental Figure 3). Neither host genetic group nor its interaction with site was significant, regardless of the order of terms were added to the model (Supplemental Table 14).

**Figure 5.**
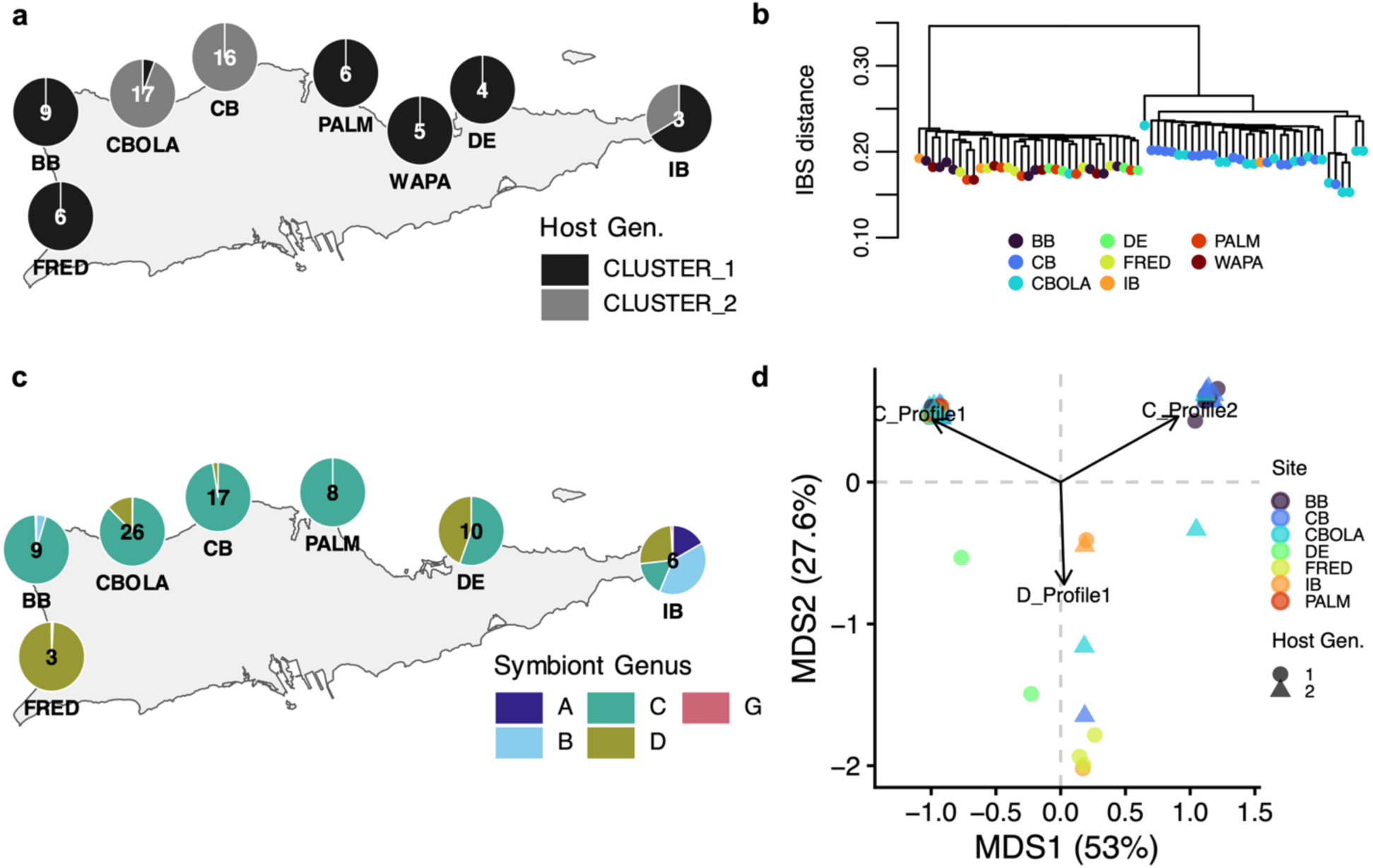
Spatial variation in *O. faveolata* host genetic structure and associated Symbiodiniaceae communities across St. Croix. (a) Relative frequencies of the two host genetic groups at each reef site. (b) Average-linkage hierarchical clustering of pairwise identity-by-state (IBS) genetic distances among host colonies; terminal points are colored by sampling site. (c) Mean relative abundances of host-associated Symbiodiniaceae genera at each site (A, *Symbiodinium*; B, *Breviolum*; C, *Cladocopium*; D, *Durusdinium*; G, *Gerakladium*). Numbers within pies in panels (a) and (c) indicate the number of samples contributing to each site-level summary. (d) Principal components analysis of SymPortal ITS2 type-profile relative abundances. Points represent individual colonies, with color indicating sampling site and shape indicating host genetic group. A small (0.05) jitter was applied to points to aid in visualization. Arrows identify the three ITS2 type profiles with the strongest loadings on the displayed axes; see Supplemental Figure 2 for all loadings and full profile strings.

Because environmental measurements were collected at the site rather than colony level, we used Bayesian hierarchical models to evaluate whether environmental similarity among reefs improved predictions of host-associated Symbiodiniaceae composition. One model included independent random intercepts for site, whereas the other allowed site effects to covary according to environmental similarity. Leave-one-out cross-validation showed equivalent predictive performance between the environmentally structured and unstructured models (ΔELPD = 0.0, SE = 2.0), providing no evidence that the environmental covariance structure improved colony-level predictions beyond site identity alone. However, Procrustes analysis found significant concordance between environmental distances and site centroid distances representing mean Symbiodiniaceae composition (Procrustes sum of squares = 0.3842, *r* = 0.7847, *p* = 0.031). Thus, abiotic conditions (e.g., temperature, turbidity) were associated with average community differences among sites, although incorporating the measured environmental similarity between sites did not improve predictions for individual colonies.

Finally, the unstructured site Bayesian model broadly supported the PERMANOVA results. Site-level variation was substantial along all three PCoA axes (Supplemental Tables 15-16), whereas neither depth nor host genetic group had a clear association with any axis.

## Discussion

By pairing repeated sampling of sediments and adult *O. faveolata* at two reefs with an island-wide coral survey, we examined whether environmental and host-associated communities responded to the same spatial and temporal factors. Reef identity provided a common thread across datasets, while the effects of time, depth and host genetic structure differed among community components. These contrasts allowed us to separate broad reef-level patterns from processes which may be specific to sediment microbiomes, environmental symbiont pools and coral hosts.

In reef sediments, taxonomic and functional configurations were strongly correlated, suggesting that microbial turnover between CB and DE extended to functional potential. Functional composition also appeared more variable at DE, whereas CB communities showed more consistent occurrence of functions related to phototrophy, environmental sensing and organic-matter processing. These patterns may reflect ecological differences between the deeper and more coral-diverse CB, and the shallower, turbid, turf-laden DE. This aligns with prior work showing that benthic composition and associated organic-matter inputs can shape microbial metabolism and functional potential across reef habitats (Dinsdale et al., 2008; Haas et al., 2011; Nelson et al., 2023; Wegley Kelly et al., 2014). Further, the high diversity of benthic cover at DE may have contributed to greater template complexity or co-extracted inhibitors, potentially reducing the consistency of ITS2 amplification from this site.

Beyond these spatial differences, both taxonomic and functional microbiome composition showed site-specific temporal change, consistent with evidence that coral reef environmental microbiomes respond to local conditions across diel and seasonal scales (Glasl et al., 2020; Wegley Kelly et al., 2019). The sampling window encompassed observed *O. faveolata* spawning and increasing sea-surface temperature, both of which could alter organic-matter inputs and microbial activity (Ceh et al., 2012; Eyre et al., 2008; Patten et al., 2008; Wild et al., 2008), although our design cannot attribute the observed turnover directly to either event. No individual KO varied among sampling days after multiple-testing correction, suggesting that temporal functional turnover was distributed across many genes rather than driven by a small set of functions. Future studies could determine whether microbial functions and taxa detected in the environment are also active in adult corals and recruits, which our host-associated lcWGS data lacked the resolution to assess.

We further examined environmental Symbiodiniaceae communities alongside those associated with nearby *O. faveolata* colonies. In both habitats, site was the primary driver of community composition. Sediment communities also showed modest, site-specific turnover after spawning, particularly at DE, whereas host-associated communities remained remarkably stable through time, consistent with the well-documented stability of adult coral–Symbiodiniaceae associations (Goulet, 2006; Thornhill et al., 2006). The stronger sediment community turnover we observed at DE may have been related to concurrent spawning of *Pseudodiploria strigosa* and *Pseudodiploria clivosa*, which are densely concentrated at the site. Further, it is possible that the enrichment of *Durusdinium* sp. ASVs at DE post-spawning may have resulted from the shedding of Symbiodiniaceae from these nearby heterospecific colonies (e.g., Hagedorn et al., 2015) which we did not measure in this study. Regardless, the strong site effect and substantial variation among sediment subsites suggest that environmental symbiont pools are structured at both reef-wide and highly localized scales.

Although environmental Symbiodiniaceae remain much less characterized than their in-host counterparts (Bell & Quigley, 2025), spatial and temporal heterogeneity may be particularly consequential during symbiosis establishment. Many corals acquire symbionts horizontally after settlement, and sediment exposure can affect both symbiont acquisition and larval performance (Adams et al., 2009; Serrano et al., 2024). Variation in the symbiont pool encountered by recruits may therefore contribute to the site-specific communities observed in adults (Scott et al., 2024). Thus, adult coral symbiont communities may represent a snapshot of the Symbiodiniaceae which were available during their settlement on the reef in years prior. However, sediment and host samples shared few ASVs, indicating that bulk sediment composition may not directly represent the symbiont pool available to *O. faveolata*. Host-compatible types may instead be rare, patchily distributed, concentrated in other environmental reservoirs, or may be more available when recruits settle. Sampling from spawning through settlement and early life history stages would help determine which environmental symbionts are acquired and whether they leave a persistent signature on adult communities.

We next expanded beyond CB and DE using the island-wide survey to ask whether site-specific differences in Symbiodiniaceae composition reflected cryptic host genetic structure rather than reef environment. Because *O. faveolata* genetic groups segregate among reefs (Cabacungan et al., 2025), consistent host-symbiont associations could produce apparent site effects. However, host genetic group did not predict ITS2 composition once the sampling site was accounted for. This was unexpected because host lineage can strongly structure symbiont associations in other coral species (Aichelman et al., 2025; Grupstra et al., 2024; Johnston et al., 2022; Scott et al., 2025). One possible explanation is transmission mode, as much of this evidence comes from vertically transmitting corals, in which inherited symbionts may remain more tightly associated with particular host lineages. Because *O. faveolata* instead acquires its symbionts horizontally, the marked differences between sediment communities at CB and DE suggest that environmental filtering may structure the available symbiont pool before host acquisition. Site-specific host communities may therefore arise not only through selection by the host, but also because recruits encounter fundamentally different symbiont pools on the reef.

Although incorporating environmental similarity did not improve colony-level predictions beyond site identity, environmental differences among reefs were correlated with their mean Symbiodiniaceae composition. Because environmental conditions were interpolated from monitoring stations rather than measured at each colony, these estimates may capture broad reef-scale differences while missing fine-scale heterogeneity. The significant site-by-depth interaction suggests that depth captured some of this finer-scale variation, consistent with depth-related differences reported within other reefs (Bongaerts et al., 2015; Eckert et al., 2020). However, depth likely represents unmeasured environmental conditions, such as light, and paired *in situ* measurements would be needed to further identify the drivers of within-reef variation.

Together, our results suggest that site captures a suite of environmental and ecological differences that shape both environmental and host-associated microbial communities. Differences in sediment microbial taxa were accompanied by differences in functional potential, suggesting that reefs differ not only in which microbes are present but likely also in the resultant microbially-driven biochemical environment. Parallel site-specific differences in environmental and coral-host Symbiodiniaceae community composition, even after accounting for host genetics, further support a role of the local environment, potentially by shaping symbiont availability before acquisition. Future work pairing *in situ* abiotic measurements with environmental and host-associated microbial sampling from spawning through recruitment is needed to provide a more complete picture of how coral symbioses become established across the coral life cycle. More broadly, our findings highlight the importance of considering the environmental microbial pool alongside host biology in understanding how coral symbioses are structured in nature.

## Data Availability

Raw read data can be found under BioProject accessions PRJNA1122865 (spatial surveys) and PRJNA1281116 (temporal surveys). All scripts, intermediate data products, and figure-generating code can be found at https://github.com/cb-scott/StCroix_SedimentAnalysis and will be archived on Zenodo upon acceptance.

## Acknowledgements

Logistical support for this project was provided by The Nature Conservancy’s USVI Coral Innovation Hub. Sampling was conducted in accordance with the U.S. Virgin Islands Department of Planning and Natural Resources (The Nature Conservancy Coral Restoration Permit DFW20052X). All bioinformatic processing was run on the Texas Advanced Computing Cluster. This study was funded by the NatureNet Science Fellowship from The Nature Conservancy to KLB, by the National Fish and Wildlife Foundation Grant 0318.20.069532 awarded to The Nature Conservancy, NSF Grant OCE-2318775 to MVM, Human Frontiers Science Program Grant 201903079001 to MVM, a grant from the Integrative Biology Department at the University of Texas at Austin to CBS, and a grant from the International Women’s Fishing Association to CBS. The funders played no role in the design or analysis of this study.

## Author Contributions

Conceptualization: CBS and KLB; Data curation: CBS, MRC and KLB; Formal analysis: CBS and MRC; Funding acquisition: CBS, MVM and KLB; Investigation: CBS, MVM, MRC, KLB, ENN, AKH and DMF; Methodology: MVM and AKH; Project administration: CBS, MVM, KLB, ENN and DMF; Resources: MVM and KLB; Software: CBS; Supervision: CBS, MVM and KLB; Visualization: CBS and MRC; Writing - Original Draft: CBS; Writing - Review & Editing: CBS, MVM, MRC, KLB, ENN, AKH and DMF

## Supplemental Tables

**Supplemental Table 1.** Spatial extent and sequencing methodologies for data included in this study. Site abbreviations correspond to Figure 1a.

| Dataset | Sites | Samples | Sequencing | Source | Accessions |
| --- | --- | --- | --- | --- | --- |
| Spatial *O. faveolata* host genetics | N = 8 (FRED, BB, CBOLA, PALM, WAPA, IB, CB, DE) | 66 | 2bRAD | Cabacungan et al., 2025; This Study | PRJNA1122865 |
| Spatial *O. faveolata* Symbiodiniaceae communities | N = 7 (FRED, BB, CBOLA, PALM, IB, CB, DE) | 80 | Amplicon/ITS2 | This Study | PRJNA1122865 |
| Temporal *O. faveolata* Symbiodiniaceae communities | N = 2 (CB, DE) | 27 | Amplicon/ITS2 | This Study | PRJNA1281116 |
| Temporal marine sediment Symbiodiniaceae communities | N = 2 (CB, DE) | 61 | Amplicon/ITS2 | This Study | PRJNA1281116 |
| Temporal marine sediment metagenomic communities | N = 2 (CB, DE) | 55 | lcWGS | This Study | PRJNA1281116 |

**Supplemental Table 2.** PERMANOVA results testing the effects of reef site (CB and DE), sampling day, depth, and their interactions on reef sediment metagenomic community composition.

| *Term* | *df* | *Sum of squares* | *R*<sup>2</sup> | *F* | *p* |
| --- | --- | --- | --- | --- | --- |
| *Site* | 1 | 0.09913 | 0.07589 | 5.0562 | **0.004** |
| *Depth* | 1 | 0.02092 | 0.01602 | 1.0670 | 0.333 |
| *Sampling Day* | 1 | 0.05560 | 0.04257 | 2.8361 | **0.030** |
| *Site × Depth* | 1 | 0.03376 | 0.02585 | 1.7220 | 0.107 |
| *Site × Sampling Day* | 1 | 0.05767 | 0.04415 | 2.9414 | **0.028** |
| *Residual* | 53 | 1.03907 | 0.79552 | — | — |
| *Total* | 58 | 1.30614 | 1.00000 | — | — |

**Supplemental Table 3.** Bayesian model results for the first three principal coordinate axes (PCoA1–PCoA3) of reef sediment metagenomic community composition, jointly modeled as responses to reef site, sampling day, and their interaction. CB was the reference site, such that site effects represent differences between DE and CB. Estimates are posterior means with standard errors (SE) and 95% credible intervals (CrI). R̂ and bulk and tail effective sample sizes (ESS) are reported as diagnostics of model convergence and sampling efficiency.

| *Response* | *Term* | *Estimate* | *SE* | *95% CrI* | $\hat{R}$ | *Bulk ESS* | *Tail ESS* |
| --- | --- | --- | --- | --- | --- | --- | --- |
| *PCoA1* | Intercept | −0.02 | 0.03 | [−0.08, 0.04] | 1.00 | 6,037 | 6,390 |
| *PCoA1* | Site: DE | 0.04 | 0.04 | [−0.05, 0.13] | 1.00 | 5,695 | 6,133 |
| *PCoA1* | Sampling Day | −0.00 | 0.00 | [−0.01, 0.00] | 1.00 | 11,867 | 9,381 |
| *PCoA1* | DE × Sampling Day | 0.01 | 0.00 | [0.00, 0.02] | 1.00 | 11,163 | 9,615 |
| *PCoA2* | Intercept | −0.04 | 0.01 | [−0.06, −0.01] | 1.00 | 6,726 | 7,474 |
| *PCoA2* | Site: DE | 0.07 | 0.02 | [0.04, 0.10] | 1.00 | 6,664 | 7,061 |
| *PCoA2* | Sampling Day | −0.00 | 0.00 | [−0.00, 0.00] | 1.00 | 16,934 | 10,684 |
| *PCoA2* | DE × Sampling Day | −0.00 | 0.00 | [−0.00, 0.00] | 1.00 | 17,867 | 9,698 |
| *PCoA3* | Intercept | 0.01 | 0.01 | [−0.01, 0.03] | 1.00 | 10,190 | 8,345 |
| *PCoA3* | Site: DE | −0.02 | 0.01 | [−0.05, 0.01] | 1.00 | 10,527 | 8,900 |
| *PCoA3* | Sampling Day | 0.00 | 0.00 | [−0.00, 0.00] | 1.00 | 15,966 | 10,007 |
| *PCoA3* | DE × Sampling Day | 0.00 | 0.00 | [−0.00, 0.01] | 1.00 | 15,610 | 9,534 |

**Supplemental Table 4.** Posterior estimates of among-subsite variation in the random intercepts for the first three principal coordinate axes (PCoA1–PCoA3) of reef sediment metagenomic taxonomic composition. Estimates represent the standard deviation (SD) among repeatedly sampled sediment subsites, with posterior standard errors (SE), 95% credible intervals (CrI), R̂, and bulk and tail effective sample sizes (ESS) reported.

| *Parameter* | *Estimate* | *SE* | *95% CrI* | *$\hat{R}$* | *Bulk ESS* | *Tail ESS* |
| --- | --- | --- | --- | --- | --- | --- |
| *PCoA1 intercept SD* | 0.06 | 0.03 | [0.01, 0.12] | 1.00 | 1,950 | 2,054 |
| *PCoA2 intercept SD* | 0.02 | 0.01 | [0.00, 0.04] | 1.00 | 2,601 | 2,570 |
| *PCoA3 intercept SD* | 0.01 | 0.01 | [0.00, 0.03] | 1.00 | 4,181 | 5,636 |

**Supplemental Table 5.** PERMANOVA results testing the effects of reef site (CB and DE), sampling day, depth, site × sampling day, and site × depth on reef sediment functional composition. Functional composition was represented by Jaccard distances calculated from KEGG Ortholog (KO) presence–absence profiles.

| *Term* | *df* | *Sum of squares* | *R*<sup>2</sup> | *F* | *p* |
| --- | --- | --- | --- | --- | --- |
| *Site* | 1 | 0.09509 | 0.13140 | 8.5452 | **0.001** |
| *Sampling Day* | 1 | 0.01780 | 0.02459 | 1.5991 | 0.091 |
| *Depth* | 1 | 0.00774 | 0.01070 | 0.6955 | 0.809 |
| *Site × Sampling Day* | 1 | 0.02946 | 0.04071 | 2.6472 | **0.014** |
| *Site × Depth* | 1 | 0.02830 | 0.03911 | 2.5434 | **0.017** |
| *Residual* | 49 | 0.54527 | 0.75349 | — | — |
| *Total* | 54 | 0.72366 | 1.00000 | — | — |

**Supplemental Table 6.**
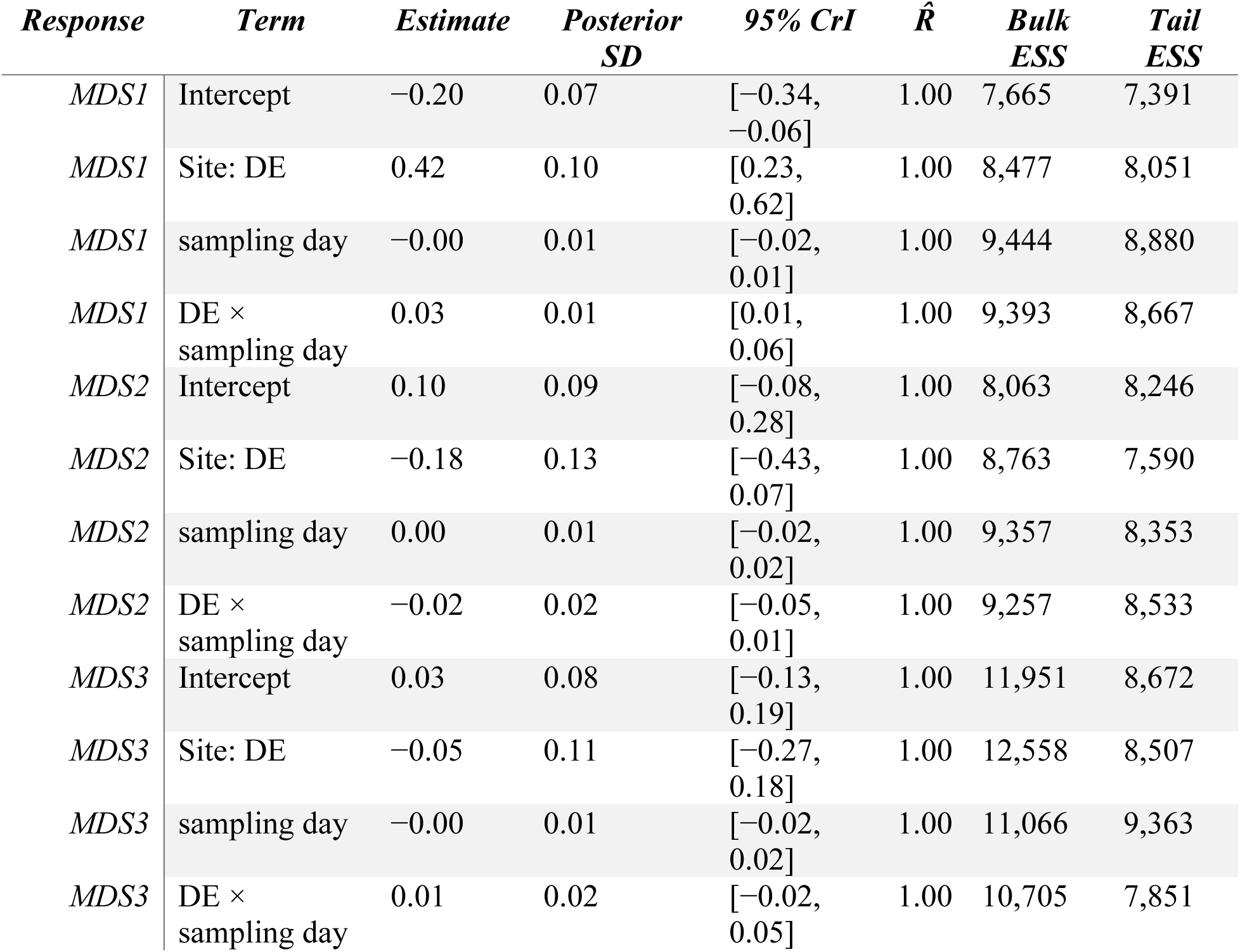
Bayesian model results for the first three principal coordinate axes (MDS1–MDS3) of reef sediment functional composition, jointly modeled as responses to reef site, sampling day, and their interaction, with sampling subsite included as a random intercept. Functional composition was represented by Jaccard distances among KO presence–absence profiles. CB was the reference site, such that site effects represent differences between DE and CB. Estimates are posterior means with posterior standard deviations and 95% credible intervals (CrI). R̂ and bulk and tail effective sample sizes (ESS) are reported as model diagnostics.

**Supplemental Table 7.** Posterior estimates of among-subsite variation in the random intercepts for the first three principal coordinate axes (MDS1–MDS3) of reef sediment functional composition. Estimates represent the standard deviation (SD) among repeatedly sampled sediment subsites, with posterior standard deviations, 95% credible intervals (CrI), R̂, and bulk and tail effective sample sizes (ESS) reported.

| *Parameter* | *Estimate* | *Posterior SD* | *95% CrI* | *$\hat{R}$* | *Bulk ESS* | *Tail ESS* |
| --- | --- | --- | --- | --- | --- | --- |
| *MDS1 intercept SD* | 0.10 | 0.06 | [0.01, 0.24] | 1.00 | 3,060 | 4,750 |
| *MDS2 intercept SD* | 0.14 | 0.08 | [0.01, 0.32] | 1.00 | 2,674 | 3,381 |
| *MDS3 intercept SD* | 0.07 | 0.06 | [0.00, 0.22] | 1.00 | 5,828 | 6,332 |

**Supplemental Table 8.** PERMANOVA results testing the effects of reef site (CB and DE), sampling day, depth, site × depth, and site × sampling day on sediment-associated *Symbiodiniaceae* community composition. Community composition was represented by Euclidean distances among centered-log-ratio-transformed ASV-group abundances.

| *Term* | *df* | *Sum of squares* | *R*<sup>2</sup> | *F* | *p* |
| --- | --- | --- | --- | --- | --- |
| *Site* | 1 | 851.46 | 0.40039 | 19.2557 | **0.001** |
| *Depth* | 1 | 81.99 | 0.03856 | 1.8543 | 0.111 |
| *Sampling day* | 1 | 42.84 | 0.02014 | 0.9688 | 0.368 |
| *Site × Depth* | 1 | 122.08 | 0.05740 | 2.7608 | **0.031** |
| *Site × Sampling day* | 1 | 99.62 | 0.04685 | 2.2530 | 0.078 |
| *Residual* | 21 | 928.59 | 0.43666 | — | — |
| *Total* | 26 | 2,126.58 | 1.00000 | — | — |

**Supplemental Table 9.**
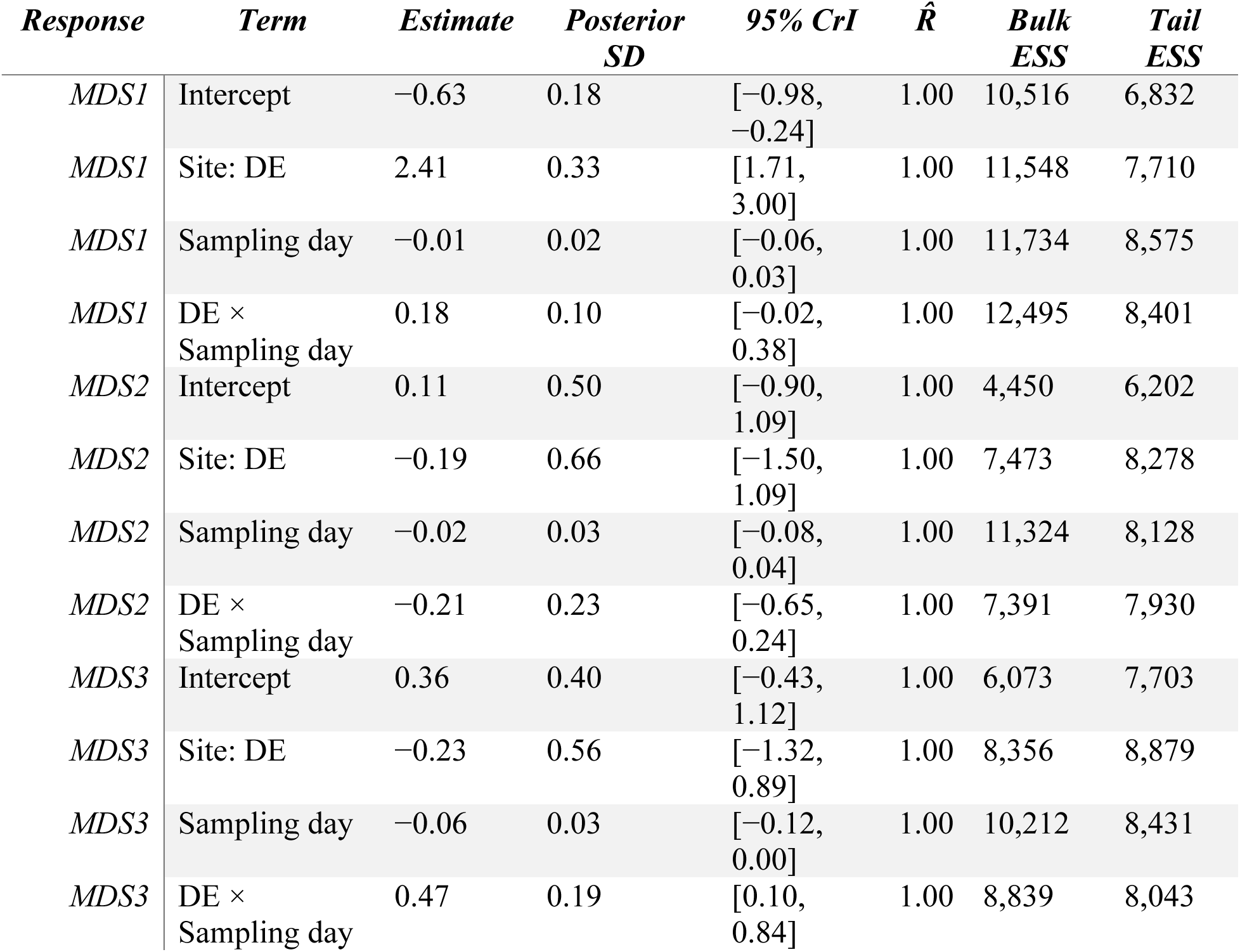
Bayesian model results for the first three principal coordinate axes (MDS1–MDS3) of sediment-associated *Symbiodiniaceae* community composition, jointly modeled as responses to reef site, sampling day, and their interaction, with sampling subsite included as a random intercept. CB was the reference site, such that site effects represent differences between DE and CB. Estimates are posterior means with posterior standard deviations and 95% credible intervals (CrI). R̂ and bulk and tail effective sample sizes (ESS) are reported as model diagnostics.

| *Response* | *Term* | *Estimate* | *Posterior*<br>*SD* | *95% CrI* | $\hat{R}$ | *Bulk*<br>*ESS* | *Tail*<br>*ESS* |
| --- | --- | --- | --- | --- | --- | --- | --- |
| *MDS1* | Intercept | −0.63 | 0.18 | [−0.98, −0.24] | 1.00 | 10,516 | 6,832 |
| *MDS1* | Site: DE | 2.41 | 0.33 | [1.71, 3.00] | 1.00 | 11,548 | 7,710 |
| *MDS1* | Sampling day | −0.01 | 0.02 | [−0.06, 0.03] | 1.00 | 11,734 | 8,575 |
| *MDS1* | DE ×<br>Sampling day | 0.18 | 0.10 | [−0.02, 0.38] | 1.00 | 12,495 | 8,401 |
| *MDS2* | Intercept | 0.11 | 0.50 | [−0.90, 1.09] | 1.00 | 4,450 | 6,202 |
| *MDS2* | Site: DE | −0.19 | 0.66 | [−1.50, 1.09] | 1.00 | 7,473 | 8,278 |
| *MDS2* | Sampling day | −0.02 | 0.03 | [−0.08, 0.04] | 1.00 | 11,324 | 8,128 |
| *MDS2* | DE ×<br>Sampling day | −0.21 | 0.23 | [−0.65, 0.24] | 1.00 | 7,391 | 7,930 |
| *MDS3* | Intercept | 0.36 | 0.40 | [−0.43, 1.12] | 1.00 | 6,073 | 7,703 |
| *MDS3* | Site: DE | −0.23 | 0.56 | [−1.32, 0.89] | 1.00 | 8,356 | 8,879 |
| *MDS3* | Sampling day | −0.06 | 0.03 | [−0.12, 0.00] | 1.00 | 10,212 | 8,431 |
| *MDS3* | DE ×<br>Sampling day | 0.47 | 0.19 | [0.10, 0.84] | 1.00 | 8,839 | 8,043 |

**Supplemental Table 10.** Posterior estimates of among-subsite variation in the random intercepts for the first three principal coordinate axes (MDS1–MDS3) of sediment-associated *Symbiodiniaceae* community composition. Estimates represent the standard deviation (SD) among repeatedly sampled sediment subsites, with posterior standard deviations, 95% credible intervals (CrI), R̂, and bulk and tail effective sample sizes (ESS) reported.

| *Parameter* | *Estimate* | *Posterior SD* | *95% CrI* | *$\hat{R}$* | *Bulk ESS* | *Tail ESS* |
| --- | --- | --- | --- | --- | --- | --- |
| *MDS1 intercept SD* | 0.19 | 0.16 | [0.01, 0.60] | 1.00 | 5,372 | 5,628 |
| *MDS2 intercept SD* | 1.35 | 0.32 | [0.84, 2.09] | 1.00 | 5,310 | 6,781 |
| *MDS3 intercept SD* | 0.92 | 0.29 | [0.41, 1.57] | 1.00 | 4,024 | 3,442 |

**Supplemental Table 11.**
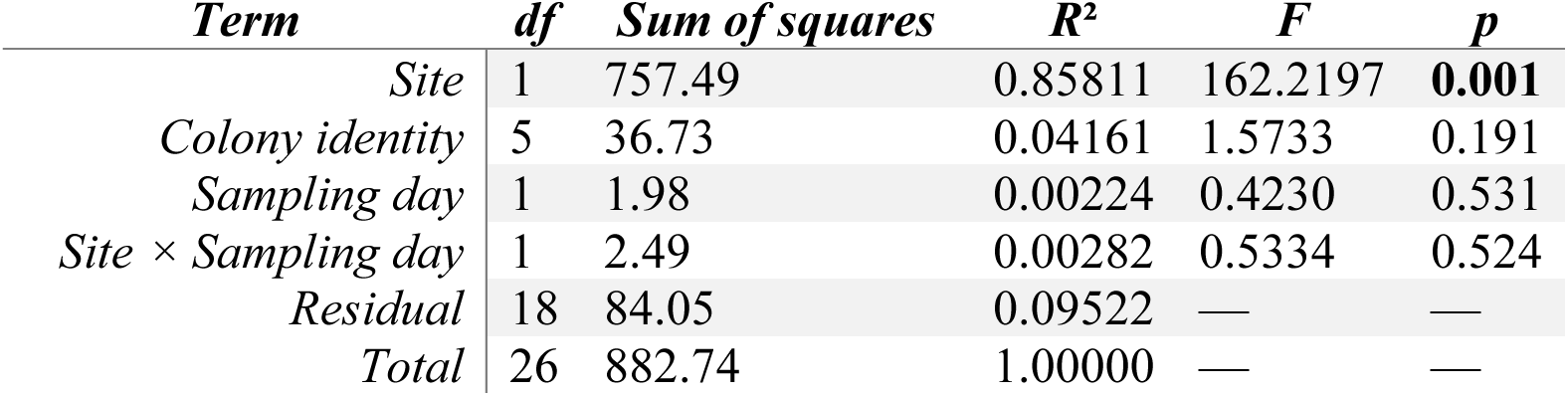
PERMANOVA results testing variation in temporally sampled host-associated *Symbiodiniaceae* community composition with reef site (CB and DE), colony identity, sampling day, and the site × sampling day interaction. Community composition was represented by Euclidean distances among centered-log-ratio-transformed ASV-group abundances.

| *Term* | *df* | *Sum of squares* | *R*<sup>2</sup> | *F* | *p* |
| --- | --- | --- | --- | --- | --- |
| *Site* | 1 | 757.49 | 0.85811 | 162.2197 | **0.001** |
| *Colony identity* | 5 | 36.73 | 0.04161 | 1.5733 | 0.191 |
| *Sampling day* | 1 | 1.98 | 0.00224 | 0.4230 | 0.531 |
| *Site × Sampling day* | 1 | 2.49 | 0.00282 | 0.5334 | 0.524 |
| *Residual* | 18 | 84.05 | 0.09522 | — | — |
| *Total* | 26 | 882.74 | 1.00000 | — | — |

**Supplemental Table 12.** Bayesian model results for the first three principal coordinate axes (MDS1–MDS3) of temporally sampled host-associated *Symbiodiniaceae* community composition, jointly modeled as responses to reef site, sampling day, and their interaction, with coral colony genotype included as a random intercept to account for repeated sampling. CB was the reference site, such that site effects represent differences between DE and CB. Estimates are posterior means with posterior standard deviations and 95% credible intervals (CrI). R̂ and bulk and tail effective sample sizes (ESS) are reported as model diagnostics.

| *Response* | *Term* | *Estimate* | *Posterior*<br>*SD* | *95%CrI* | $\hat{R}$ | *Bulk*<br>*ESS* | *Tail*<br>*ESS* |
| --- | --- | --- | --- | --- | --- | --- | --- |
| *MDS1* | Intercept | −0.98 | 0.11 | [−1.18, −0.76] | 1.00 | 6,863 | 5,027 |
| *MDS1* | Site: DE | 2.02 | 0.16 | [1.70, 2.29] | 1.00 | 5,629 | 4,028 |
| *MDS1* | sampling day | 0.01 | 0.01 | [−0.01, 0.03] | 1.00 | 6,953 | 8,277 |
| *MDS1* | DE ×<br>sampling day | −0.01 | 0.01 | [−0.04, 0.02] | 1.00 | 7,052 | 7,532 |
| *MDS2* | Intercept | 0.07 | 0.44 | [−0.81, 0.95] | 1.00 | 8,066 | 7,608 |
| *MDS2* | Site: DE | −0.08 | 0.56 | [−1.20, 1.02] | 1.00 | 8,542 | 7,913 |
| *MDS2* | sampling day | −0.01 | 0.03 | [−0.08, 0.05] | 1.00 | 9,003 | 7,874 |
| *MDS2* | DE ×<br>sampling day | 0.01 | 0.06 | [−0.10, 0.12] | 1.00 | 8,689 | 7,787 |
| *MDS3* | Intercept | 0.08 | 0.38 | [−0.67, 0.82] | 1.00 | 8,210 | 8,365 |
| *MDS3* | Site: DE | −0.23 | 0.50 | [−1.21, 0.76] | 1.00 | 8,995 | 8,551 |
| *MDS3* | sampling day | 0.02 | 0.03 | [−0.04, 0.09] | 1.00 | 6,915 | 7,478 |
| *MDS3* | DE ×<br>sampling day | −0.06 | 0.06 | [−0.17, 0.06] | 1.00 | 7,027 | 8,152 |

**Supplemental Table 13.**
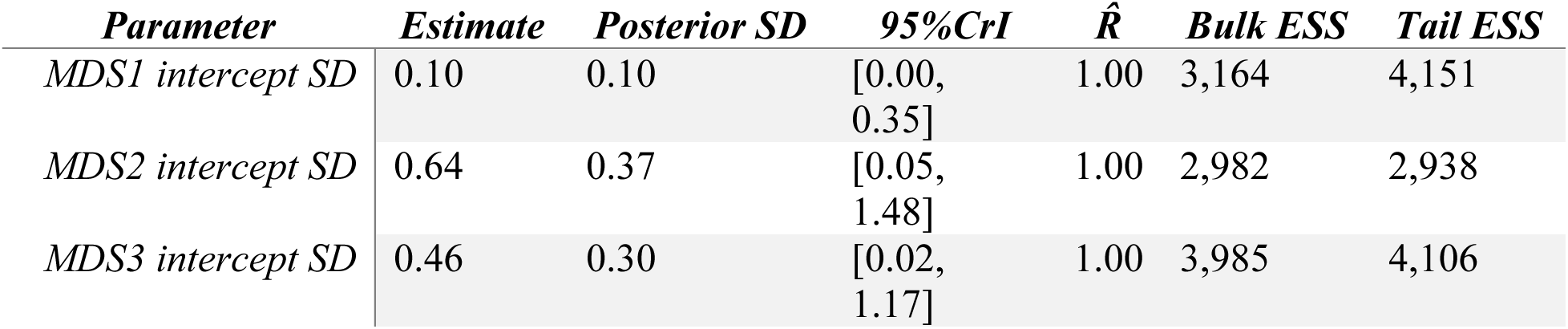
Posterior estimates of among-colony variation in the random intercepts for the first three principal coordinate axes (MDS1–MDS3) of temporally sampled host-associated *Symbiodiniaceae* community composition. Estimates represent the standard deviation (SD) among repeatedly sampled coral colonies, with posterior standard deviations, 95% credible intervals (CrI), R̂, and bulk and tail effective sample sizes (ESS) reported.

| *Parameter* | *Estimate* | *Posterior SD* | *95%CrI* | *$\hat{R}$* | *Bulk ESS* | *Tail ESS* |
| --- | --- | --- | --- | --- | --- | --- |
| *MDS1 intercept SD* | 0.10 | 0.10 | [0.00, 0.35] | 1.00 | 3,164 | 4,151 |
| *MDS2 intercept SD* | 0.64 | 0.37 | [0.05, 1.48] | 1.00 | 2,982 | 2,938 |
| *MDS3 intercept SD* | 0.46 | 0.30 | [0.02, 1.17] | 1.00 | 3,985 | 4,106 |

**Supplemental Table 14.** PERMANOVA results testing the effects of reef site, host genetic group, depth, site × host genetic group, and site × depth on spatial variation in host-associated *Symbiodiniaceae* community composition. Community composition was represented by Bray– Curtis dissimilarities among SymPortal ITS2 type-profile relative abundances.

| *Term* | *df* | *Sum of squares* | *R*<sup>2</sup> | *F* | *p* |
| --- | --- | --- | --- | --- | --- |
| *Site* | 6 | 5.1382 | 0.31189 | 3.8370 | **0.001** |
| *Host genetic group* | 1 | 0.1277 | 0.00775 | 0.5721 | 0.642 |
| *Depth* | 1 | 0.1199 | 0.00728 | 0.5372 | 0.602 |
| *Site × Host genetic group* | 1 | 0.1273 | 0.00773 | 0.5704 | 0.617 |
| *Site × Depth* | 2 | 1.5873 | 0.09635 | 3.5560 | **0.006** |
| *Residual* | 42 | 9.3738 | 0.56900 | — | — |
| *Total* | 53 | 16.4742 | 1.00000 | — | — |

**Supplemental Table 15.**
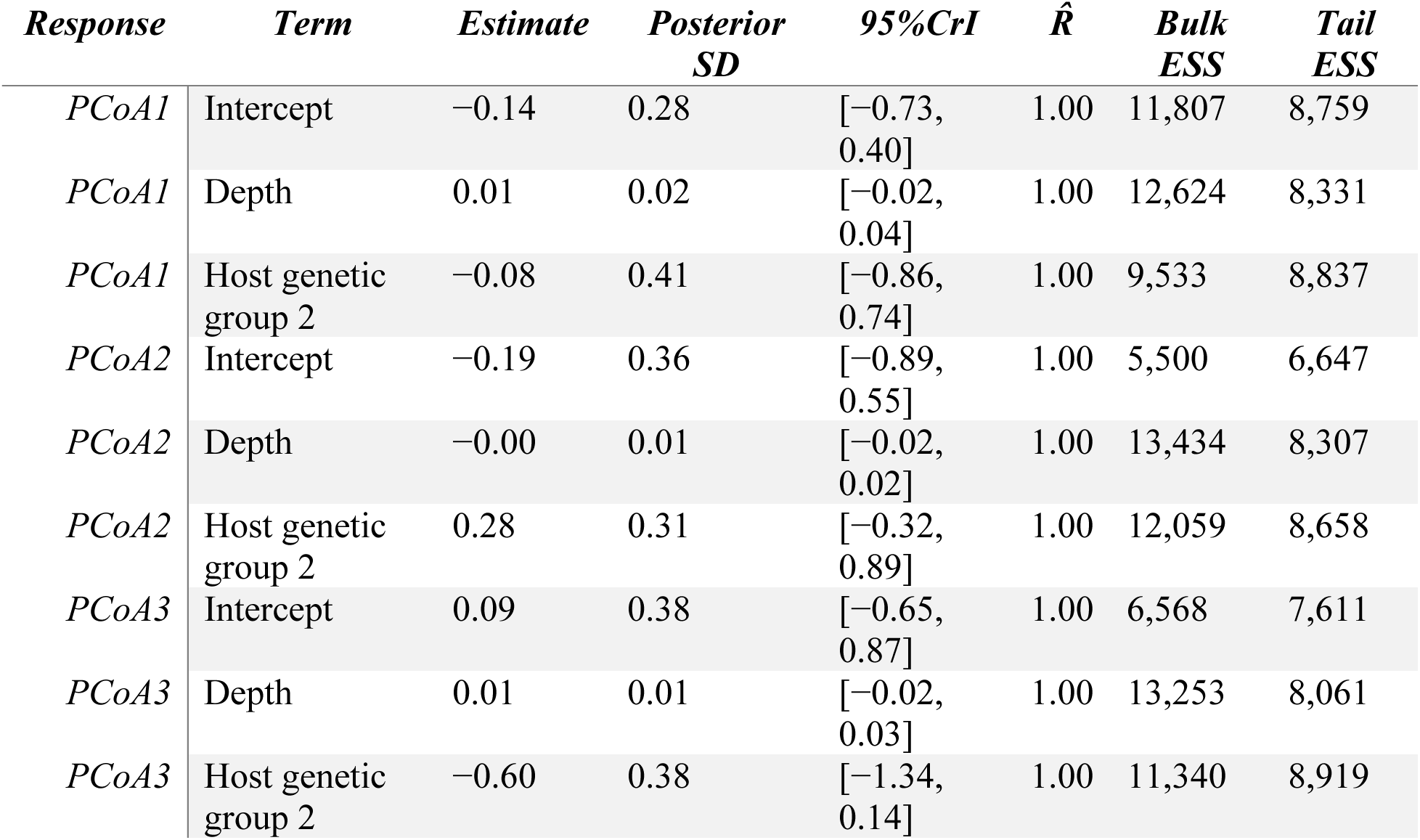
Bayesian model results for the first three principal coordinate axes (PCoA1–PCoA3) of spatial host-associated *Symbiodiniaceae* community composition. Depth and host genetic group were included as fixed effects, and reef site was included as an independent random intercept to account for clustering of colonies within sites. Host genetic group 1 was the reference group. Estimates are posterior means with posterior standard deviations and 95% credible intervals (CrI). R̂ and bulk and tail effective sample sizes (ESS) are reported as model diagnostics.

**Supplemental Table 16.** Posterior estimates of among-site variation in the random intercepts for the first three principal coordinate axes (PCoA1–PCoA3) of spatial host-associated *Symbiodiniaceae* community composition in the model with independent site effects. Estimates represent the standard deviation (SD) among reef sites, with posterior standard deviations, 95% credible intervals (CrI), R̂, and bulk and tail effective sample sizes (ESS) reported.

| *Parameter* | *Estimate* | *Posterior SD* | *95%CrI* | $\hat{R}$ | *Bulk ESS* | *Tail ESS* |
| --- | --- | --- | --- | --- | --- | --- |
| *PCoA1 intercept SD* | 0.37 | 0.25 | [0.02, 0.97] | 1.00 | 3,674 | 5,471 |
| *PCoA2 intercept SD* | 0.85 | 0.28 | [0.44, 1.52] | 1.00 | 5,091 | 7,032 |
| *PCoA3 intercept SD* | 0.84 | 0.30 | [0.40, 1.56] | 1.00 | 4,768 | 6,471 |

## Supplemental Figures

**Supplemental Figure 1.**
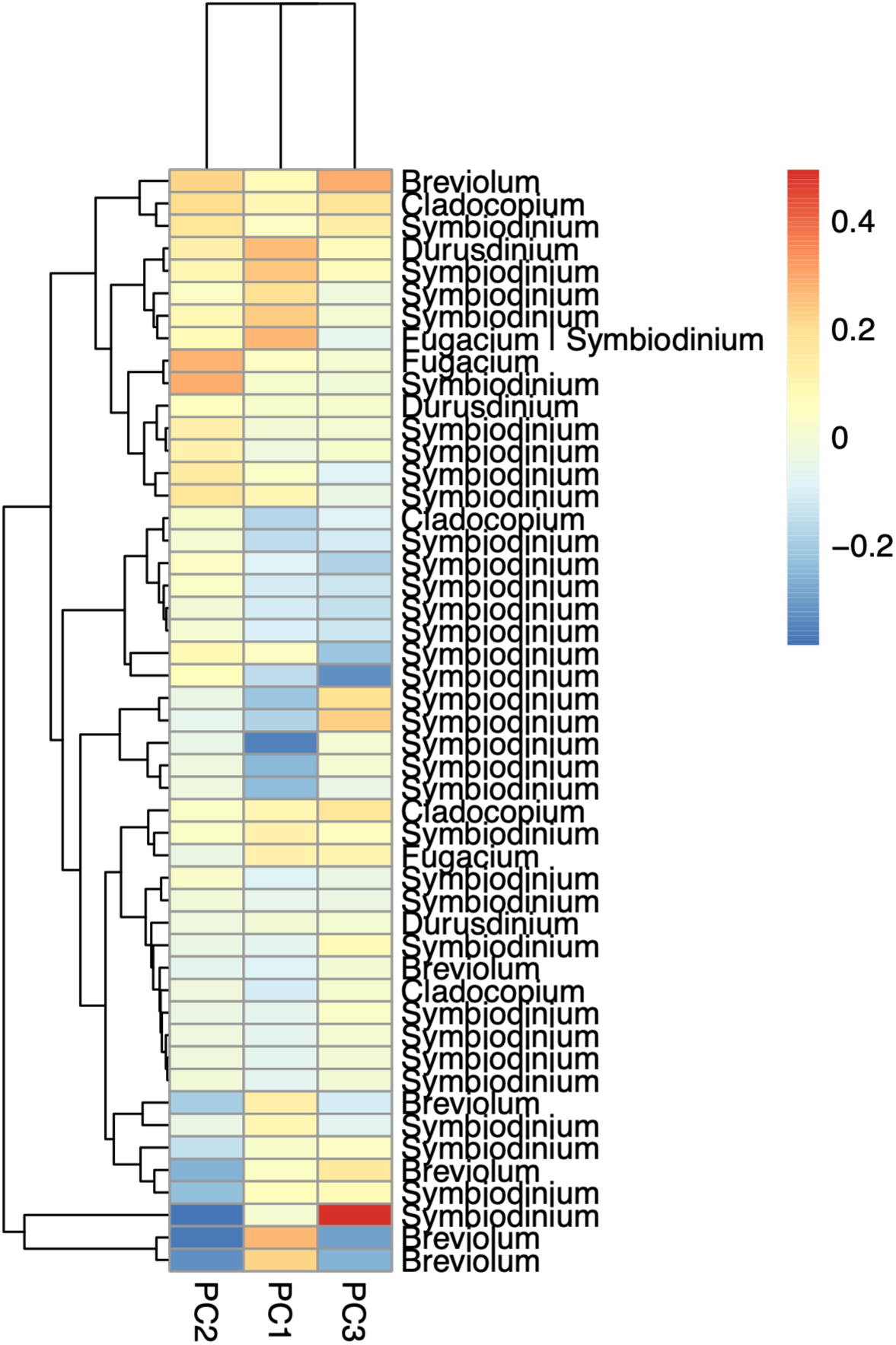
Loadings of sediment-associated *Symbiodiniaceae* ASV groups on the principal coordinate axes used to summarize variation in sediment *Symbiodiniaceae* community composition. Each row is a separate ASV group; the genus-level taxonomic assignment of that ASV group is given. Loadings indicate the direction and relative contribution of individual ASV groups to variation along each ordination axis.

**Supplemental Figure 2.**
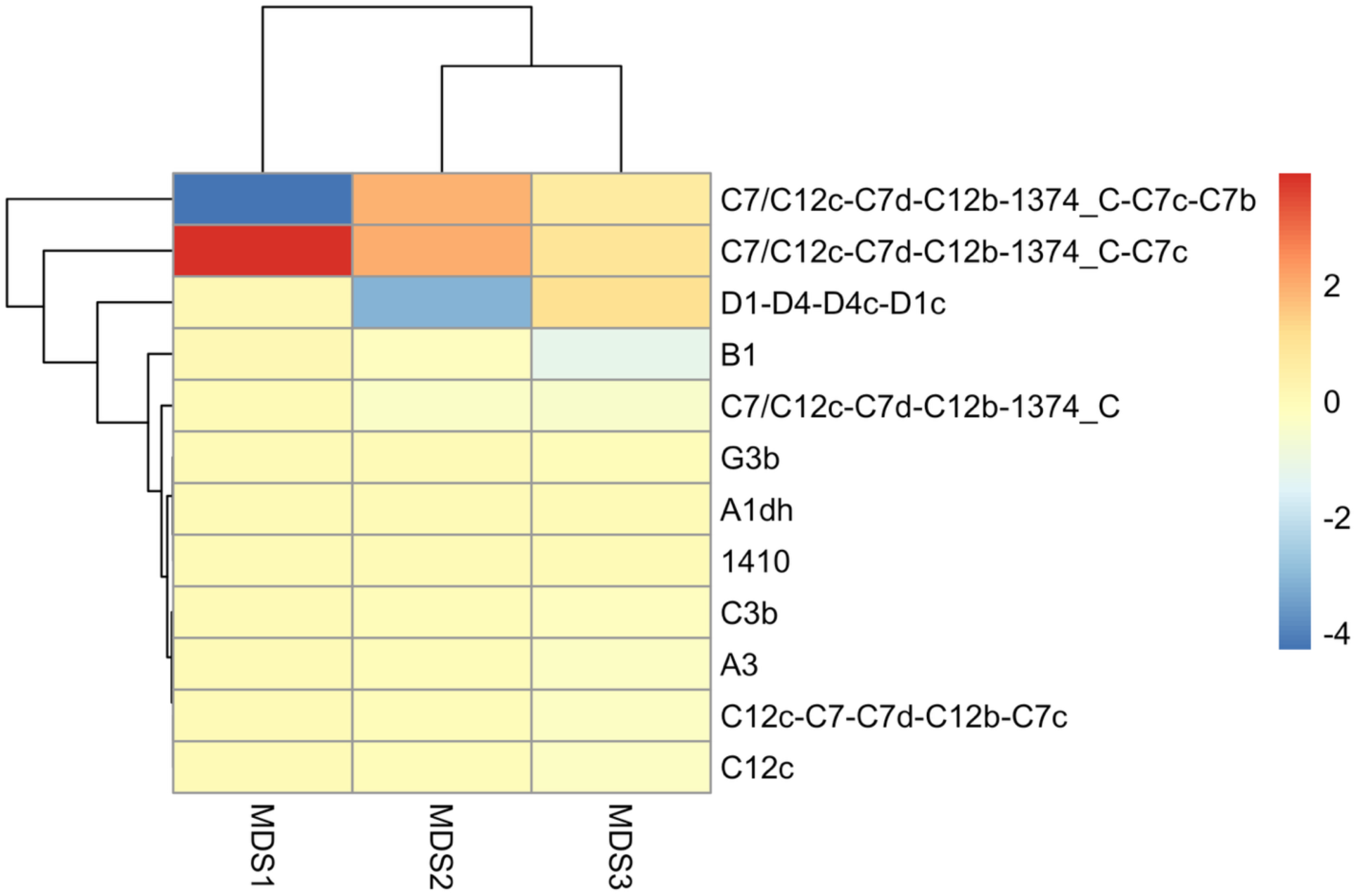
Loadings of host-associated *Symbiodiniaceae* ITS2 type profiles from Symportal on the principal coordinate axes used to summarize variation in coral-associated symbiont community composition. Loadings indicate the direction and relative contribution of individual ITS2 type profiles to variation along each ordination axis.

**Supplemental Figure 3.**
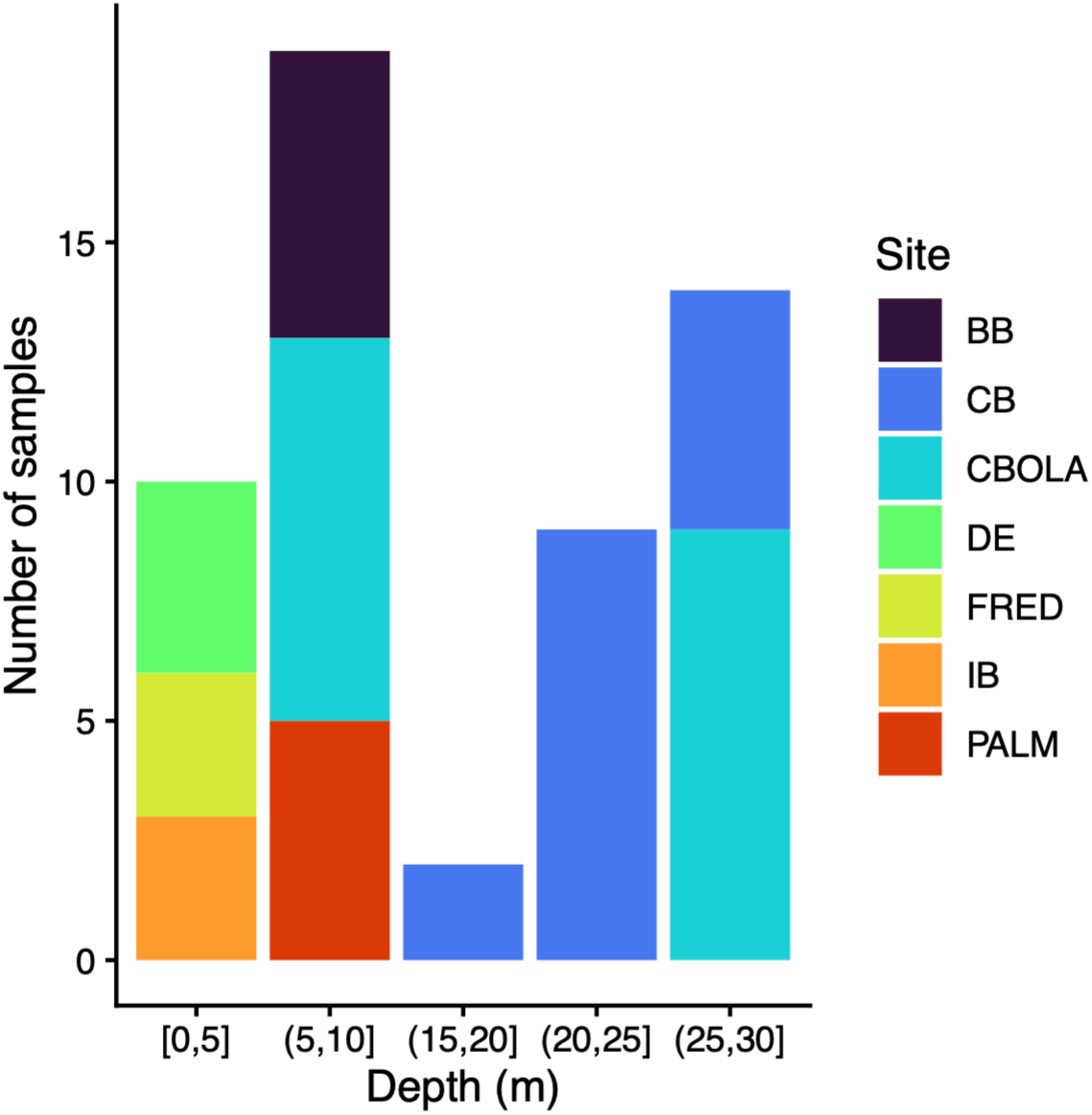
Distribution of sampling depths across reef sites included in the spatial host-associated *Symbiodiniaceae* survey. CB and CBOLA are the only sites that had >5m depth variation sampled.

