## Supplemental File 1 for "From sediment to coral symbionts: persistent spatial structure across reef microbiomes"

**Supplementary File 1:** Genomic DNA isolation protocol

First, make the 2% CTAB extraction buffer:

Ingredients for 200 ml:

2 % CTAB a.k.a. Hexadecyltrimethylammonium bromide (4.0 g)

100 mM Tris pH 8 (20 ml of 1.0 M sol)

20 mM EDTA (16 ml of 0.25 M sol)

164 ml H2O

Dissolve CTAB before adding 1.4 M NaCl (16.4 g). Stir on a hot plate with a little warmth until CTAB is dissolved.

Then add NaCl and continue stirring on hot plate until dissolved.

The following amounts are for 1 sample. Before beginning: Get ice. Place aliquot of isopropanol in the freezer. Preheat heat block to 42°C. Preheat elution buffer aliquot to 65°C.

1. In a 2mL bead beater tube, add enough beads to fill the conical bottom of the tube. Then add **1.6µL** beta merceptoethanol, **1µL** proteinase K, and **1µL** RNAse A to **800µL** of CTAB extraction buffer. Place on ice.

2. Transfer coral fragment onto a clean kimwipe using clean forceps. Cut into smaller pieces with a clean razor blade if needed. Blot away excess ethanol.

3. Add sample to the tube with beads.

4.. Macerate sample in the bead beater for 40 seconds.

5. Incubate sample at 42°C for 1 hour or overnight.

6. Spin samples in a tabletop centrifuge for 15 minutes at max speed. Transfer aqueous phase to a clean gel-lock phase tube.

7. Add **800µL** (1 volume) chloroform/isoamyl alcohol (24:1) and vortex for a few seconds. Leave on ice 1 minute. Vortex again, 1-2 seconds.

8. Spin max speed for 20 minutes at 4°C.

9. Pipette off the aqueous phase and place in a clean 1.5 ml tube.

8. Add **550µL** (2/3 volume) of ice cold isopropanol and gently mix by inverting.

9. Incubate for 20 minutes at -20°C.

10. Centrifuge at max speed for 20 minutes at 4°C.

11. Pipette off the supernatant and discard.

12. Add **1000µL** 80% ethanol. Gently wash EtOH around the tube.

13. Centrifuge for 5 minutes at 4°C, max speed.

14. Pipette off the supernatant and discard. Let the pellets air dry upside down for 15 minutes in the hood.

15. Resuspend DNA in 30µL of warm (65°C) elution buffer.

16. Nanodrop and store samples at -20°C.
