## Supplemental File 2 for "From sediment to coral symbionts: persistent spatial structure across reef microbiomes"

**Reagent List**

| Tagmentase (Tn5 Transposon)-unloaded | Diagenode C01070010-10 |
| --- | --- |
| Tagmentation Buffer 2X | Diagenode C01019043-100 |
| Adaptors A1-4 and B1-4  (resuspended in 40mM Tris-HCl (pH8.0), 50mM NaCl) | IDT DNA |
| KAPA HiFi HotStart ReadyMix | Roche 7958927001 |
| P5 and P7 primers | IDT DNA |
| Barcode (index) oligos ([same as 2bRAD barcodes](https://github.com/z0on/2bRAD_denovo)) | IDT DNA |
| Mag-Bind® TotalPure NGS | Omega Bio-Tek M1378-00 |
| 96 well magnetic plate | Thermo Fisher 12331D |

**Table 1. Reagent List**

| **Name** | **Sequence** | **Supplier** | **Purification** |
| --- | --- | --- | --- |
| Mosaicend_reverse | CTGTCTCTTATACACATCT[PHO] | IDT | 100 nmole DNA Oligo |
| Mosaicend_AdapterA | **CTACACGACGCTCTTCCGATCTAGATGTGTATAAGAGACAG** | IDT | 25 nmole DNA Oligo |
| Mosaicend_AdapterA2 | **CTACACGACGCTCTTCCGATCTGAGATGTGTATAAGAGACAG** | IDT | 25 nmole DNA Oligo |
| Mosaicend_AdapterA3 | **CTACACGACGCTCTTCCGATCTCTTAGATGTGTATAAGAGACAG** | IDT | 25 nmole DNA Oligo |
| Mosaicend_AdapterA4 | **CTACACGACGCTCTTCCGATCTTCCSAGATGTGTATAAGAGACAG** | IDT | 25 nmole DNA Oligo |
| Mosaicend_AdapterB | **CAGACGTGTGCTCTTCCGATCTAGATGTGTATAAGAGACAG** | IDT | 25 nmole DNA Oligo |
| Mosaicend_AdapterB1 | **CAGACGTGTGCTCTTCCGATCTTAGATGTGTATAAGAGACAG** | IDT | 25 nmole DNA Oligo |
| Mosaicend_AdapterB2 | **CAGACGTGTGCTCTTCCGATCTGCCAGATGTGTATAAGAGACAG** | IDT | 25 nmole DNA Oligo |
| Mosaicend_AdapterB3 | CAGACGTGTGCTCTTCCGATCT**CTTSAGATGTGTATAAGAGACAG** | IDT | 25 nmole DNA Oligo |
| ILL-BC (i7 barcode) | CAAGCAGAAGACGGCATACGAGATGCTACCGTGACTGGAGTT**CAGACGTGTGCTCTTCCGAT** | IDT | 20 nmole Ultramer DNA Oligo |
| TruSeq (i5 barcode) | AATGATACGGCGACCACCGAGATCTACACATCACGACACTCTTTCC**CTACACGACGCTCTTCCGATCT** | IDT | 20 nmole Ultramer DNA Oligo |
| P5 primer | AATGATACGGCGACCACCGA | IDT | 25 nmole DNA Oligo |
| P7 primer | CAAGCAGAAGACGGCATACGA | IDT | 25 nmole DNA Oligo |

**Table 2. Oligo Sequences of adaptors to anneal to the Tn5 transposon**

4 different versions of the two adaptors were created to improve sequencing results. Illumina reads better when there is diversity so the bases in red were added to add diversity into the sequences. The bold bases are the required sequence to anneal to the Tn5 consensus regions. The green and blue sequences are the ones that individual users can change to correspond with their specific barcoding system. See the last two rows of the table for examples of our dual index barcodes.

**Specification of adaptor design allowing flexibility in barcode usage**

Utilizing Tagmentase (Tn5 transposase) allows DNA to be sheared and adaptor sequences to be added in one step. This allows for a more efficient and cost effective process. Additionally, as Tagmentase Tn can come “unloaded”, any oligo of choice can be annealed onto it such that it works with an already established barcode system (1-3). Adaptors are designed to contain the 19 bp Tn5 mosaic ends (bolded in Table 2) recognized by the transposase and sequences complementary to our barcode system. Unlike other studies, we did not need to use any expensive kits and used our own barcodes designed for the TruSeq Illumina system and used a more cost-effective PCR reagent. The cost of our library prep is ~$2.75 versus procedure using preloaded Tn5 which is ~$6.50 per sample (4-5). Our protocol uses the hyperactive mutant of Tn5 transposase supplied unloaded from Diagenode.

**Detailed Protocol**

**Reagents to Prepare**

1. Prepare the following reagents
   a. **10 µM P5 and P7 primers**
   b. **2 µM** of the respective **TruSeq Un** oligos and **ILL-BC** oligos
      i. Need enough to create unique combination of TruSeq Un and ILL-BC for each sample
   c. **1X Tagmentation Buffer**
      i. Dilute from the 2X Tagmentation buffer using milliQ water

**Normalization**

1. Run PicoGreen Assay to obtain DNA concentrations
2. Use values to normalize samples in water to 2.5 ng/µL
   a. Range of 0.5-10 ng/µl has been tested before

**Tagmentase Assembly ([Diagenode Protocol](https://www.diagenode.com/files/protocols/PRO-Transposome-Assembly-V2.pdf))**

1. In separate PCR tubes, mix equal volumes of the **100 µM** reverse oligo with each **100 µM** adaptor A and each **100 µM** adaptor B (total of 8 mixes)
2. Mix and spin down tubes before running the following PCR cycle:

95°C for 5 minutes

65°C for 5 minutes

Hold at 4°C

3. Mix equal volume of each annealed adaptor A and annealed adaptor B into one tube
   a. This can be stored at -20°C before the Tagmentase is added
4. Add an equal volume of Tagmentase to the mixture
5. Vortex briefly and incubate at 23°C for 45 minutes
6. Dilute assembled Tamgentase to concentration of 50 ng/µL with 1X tagmentation buffer
   a. Tagmentase comes as **2,000 ng/µL** and is diluted to 1,000 ng/µL when it is mixed with the adaptors
7. Use immediately
   a. (can be stored in -20 for at least 6 months)

**Tagmentation**

1. Preheat thermocycler to 55°C
2. Thaw primers and oligos and Kapa HiFi master mix for the next step
3. Prepare the following master mix multiplying each reagent by the number of samples * 1.1

| Reagent | Volume per Sample |
| --- | --- |
| Tagmentation Buffer | 1.25 µL |
| Tagmentase Enzyme | 0.25 µL |

4. Distribute master mix in PCR strip tubes if prepping a large number of samples
5. Use a multichannel pipet to add **1.5 µL of master mix** to each well
6. Add **1 µL of DNA** to each well and mix by pipetting up and down at least 10 times
7. Seal plate/close tubes and centrifuge briefly
8. Incubate plate/tubes for 10 minutes at 55℃
9. Centrifuge briefly after incubation

**Amplification**

1. Prepare the following master mix for **each of the TruSeq Un oligos** being used by multiplying the volume by the number of samples + 0.5

| Reagent | Volume per sample |
| --- | --- |
| MilliQ or Nuclease-Free water | 3.68 µL |
| Kapa Hifi Readymix | 11.25 µL |
| 10 µM P5 | 0.56 µL |
| 10 µM P7 | 0.56 µL |
| 2 µM TruSeq Un_ | 2.8 µL |

2. Add **2.8 µL** of respective **2 µM ILL-BC_** to each sample
   a. Make sure each sample has a unique combo of TruSeq_Un and ILL-BC
   b. TruSeq is unique to each row and Ill-BC is unique to each column generally
3. Add **18.85 µL**of the respective **TruSeq Un master mix** to each sample, mix via pipetting
4. Seal the plate/close the tubes and centrifuge briefly
5. Run the following PCR program

72°C for 3 min

98°C for 2:45 min

98°C for 15sec

62°C for 30sec	x 17 cycles

72°C for 3min

72°C for 1 min

6. Check on 2% gel run at 80 V for 50 min, should see nice bright smears (see picture on next page)
   a. If preparing hundreds of samples, check a subset
7. Samples can be stored in fridge before cleaning

**Pool Samples**

1. If running lots of samples, they can be pooled at this step before cleaning

**PCR Clean-up and Size Selection**

1. Briefly centrifuge plate/tubes
2. Bring Mag-Bind® TotalPure NGS beads to room temperature and resuspend the beads via vortexing
3. Pipet **0.57X volume of beads (0.57x of PCR product volume)** into each well and pipet up and down 10 times to mix well
   a. This will size select for about 500 bp and larger
4. Incubate at room temperature for 10 minutes
5. Place plate onto magnet until the solution clears
6. Remove and discard supernatant
7. Add **150 µL 80% ethanol** to each well and let stand for **30 seconds**
8. Remove ethanol and repeat step 7
9. Remove residual ethanol
10. Let dry for no more than 5 minutes
11. Resuspend beads in **32 µL 10mM Tris, ph8**
    a. Or any elution buffer or water
12. Pipet at least 10 times to resuspend beads
13. Incubate for 5 minutes at room temperature
14. Place plate onto magnet until solution clears
15. Transfer supernatant (only **30 µL** to avoid bead contamination) to new plate
